# Contextual flexibility in *Pseudomonas aeruginosa* central carbon metabolism during growth in single carbon sources

**DOI:** 10.1101/828012

**Authors:** Stephen K. Dolan, Michael Kohlstedt, Stephen Trigg, Pedro Vallejo Ramirez, Christoph Wittmann, Clemens F. Kaminski, Martin Welch

## Abstract

*Pseudomonas aeruginosa* is an opportunistic human pathogen, particularly noted for causing infections in the lungs of people with cystic fibrosis (CF). Previous studies have shown that the gene expression profile of *P. aeruginosa* appears to converge towards a common metabolic program as the organism adapts to the CF airway environment. However, at a systems level, we still have only a limited understanding of how these transcriptional changes impact on metabolic flux. To address this, we analysed the transcriptome, proteome and fluxome of *P. aeruginosa* grown on glycerol or acetate. These carbon sources were chosen because they are the primary breakdown products of airway surfactant, phosphatidylcholine, which is known to be a major carbon source for *P. aeruginosa* in the CF airways. We show that the flux of carbon through central metabolism is radically different on each carbon source. For example, the newly-recognised EDEMP cycle plays an important role in supplying NADPH during growth on glycerol. By contrast, the EDEMP cycle is attenuated during growth on acetate, and instead, NADPH is primarily supplied by the isocitrate dehydrogenase(s)-catalyzed reaction. Perhaps more importantly, our proteomic and transcriptomic analyses reveal a global remodelling of gene expression during growth on the different carbon sources, with unanticipated impacts on aerobic denitrification, electron transport chain architecture, and the redox economy of the cell. Collectively, these data highlight the remarkable metabolic plasticity of *P. aeruginosa*; a plasticity which allows the organism to seamlessly segue between different carbon sources, maximising the energetic yield from each.

**Importance:** *Pseudomonas aeruginosa* is an opportunistic human pathogen, well-known for causing infections in the airways of people with cystic fibrosis. Although it is clear that *P. aeruginosa* is metabolically well-adapted to life in the CF lung, little is currently known about how the organism metabolises the nutrients available in the airways. In this work, we use a combination of gene expression and isotope tracer (“fluxomic”) analyses to find out exactly where the input carbon goes during growth on two CF-relevant carbon sources, acetate and glycerol (derived from the breakdown of lung surfactant). We find that carbon is routed (“fluxed”) through very different pathways during growth on these substrates, and that this is accompanied by an unexpected remodelling of the cell’s electron transfer pathways. Having access to this “blueprint” is important because the metabolism of *P. aeruginosa* is increasingly being recognised as a target for the development of much-needed antimicrobial agents.

## Introduction

*Pseudomonas aeruginosa* (Pa) is an opportunistic pathogen. This cosmopolitan microbe has become one of the most frequent causative agents of nosocomial infection [1]. It is also well-known for colonising the airways of cystic fibrosis (CF) patients; a spatially and chemically heterogenous environment characterised by gradients of oxygen and nutrients. To survive in this niche, Pa must therefore overcome numerous challenges [2,3]. Indeed, recent studies have suggested that CF-adapted *P. aeruginosa* exhibits distinct physiological adaptations, including a tailored preference for specific carbon sources, increased requirement for oxygen, and decreased fermentation [4]. Moreover, and despite extensive studies into the physiology and metabolism of *P. aeruginosa*, we still lack a clear understanding of how the growth and assimilation of carbon is controlled in this organism. Uncovering these processes is central to developing future treatment strategies.

Carbon source utilisation by *P. aeruginosa* is hierarchical, and the organism displays marked diauxy during growth on mixed carbon sources [5]. *In vitro*, the preferred carbon source of domesticated *P. aeruginosa* strains includes tricarboxylic acid cycle intermediates and amino acids. Intriguingly, and unlike many enteric bacteria, glucose is not especially favoured, even though it is present at millimolar concentrations in CF secretions [6]. Instead, adapted CF airway isolates seem to prefer glycerol during exponential growth. The precise reason for this is unclear, as glycerol *per se* is not abundant in CF sputum [4]. However, the surfactant phosphatidylcholine (PC) is abundant in the CF airways, and *P. aeruginosa* is known to secrete lipases which can cleave PC to yield phosphorylcholine, glycerol, and long-chain fatty acids (FAs) such as palmitate. The liberated glycerol is then metabolised through the action of enzymes encoded by the *glp* operon, whereas the FAs are iteratively degraded by β-oxidation to yield acetyl-CoA. The acetyl moiety is then shuttled into the TCA cycle and glyoxylate shunt to generate energy and gluconeogenic precursors for biomass production, respectively. Other sources of short chain fatty acids in the CF airways have also been recently identified [7]. Indeed, acetate derived from tracheobronchial mucin breakdown by anaerobes has been reported at concentrations in excess of 5 mM in CF sputum, and reconstitution of the CF airway microbiota in mucin-containing medium *in vitro* leads to >30 mM acetate accumulating [7]. Glycerol and acetate metabolism therefore occupy an important crossroads in *P. aeruginosa* pathophysiology [8]. Glycerol is also a preferred carbon source for alginate synthesis by CF isolates of *P. aeruginosa* and promotes the appearance of mucoid variants when present at high concentrations [9].

Despite the obvious importance of these substrates for pathophysiology, we still do not know how glycerol and acetate impact on the metabolism, redox balance, and gene expression profile of *P. aeruginosa*. Much of what we think we know has been gleaned by extrapolation from other bacterial species, yet those species often occupy very different niches compared with *P. aeruginosa*, and display different substrate preferences. To redress this, in the current work, we use a combination of ‘omics approaches (transcriptomics, proteomics, and [^13^C] fluxomics), coupled with reverse genetics, to systematically investigate the pathway(s) of carbon assimilation during growth on acetate and glycerol. Surprisingly, not only do these different carbon sources lead to a “rewiring” of central metabolism; they also give rise to profound changes in the expression of pathogenicity-associated functions including denitrification, redox balance mechanisms and the electron transport chain. These data underline the striking metabolic flexibility of *P. aeruginosa*, which allows this organism to carry out efficient, real-time free energy conservation, a trait which is likely to aid its ability to proliferate in diverse environmental conditions.

## Results

### Comparative transcriptomic, proteomic and fluxomic analysis of PAO1 cultured on glycerol or acetate as a sole carbon sources reveals global changes in central carbon metabolism

We first examined the growth characteristics of *P. aeruginosa* during cultivation on acetate and glycerol as single carbon sources. This revealed that *P. aeruginosa* grows more slowly in glycerol (growth rate, µ_max_ = 0.37±0.01 h^-1^) than it does in acetate (µ_max_ = 0.80±0.01 h^-1^), glucose (µ_max_ = 0.88±0.05 h^-1^) or succinate (µ_max_ = 0.87±0.05 h^-1^) (Figure S2). To investigate this further, we examined the transcriptome, proteome and fluxome of *P. aeruginosa* during the assimilation of glycerol and acetate. Our analysis was carried out on cells grown to mid-exponential phase (OD_600_ = 0.5) in baffled shake flasks containing MOPS-buffered media. This allowed us to elucidate specific impact of carbon source utilization on *P. aeruginosa* metabolism and physiology without the confounding factors of nutrient and oxygen limitation that accompany entry into the stationary phase.

Acetate and glycerol have different entry points into *P. aeruginosa* central carbon metabolism, and are also thought to have distinct effects on redox metabolism [10–13]. We therefore anticipated a carbon source-specific impact on the expression of enzymes (and corresponding fluxes) involved in the relevant pathways. These pathways are summarised in Figure 1.

**Figure 1.**
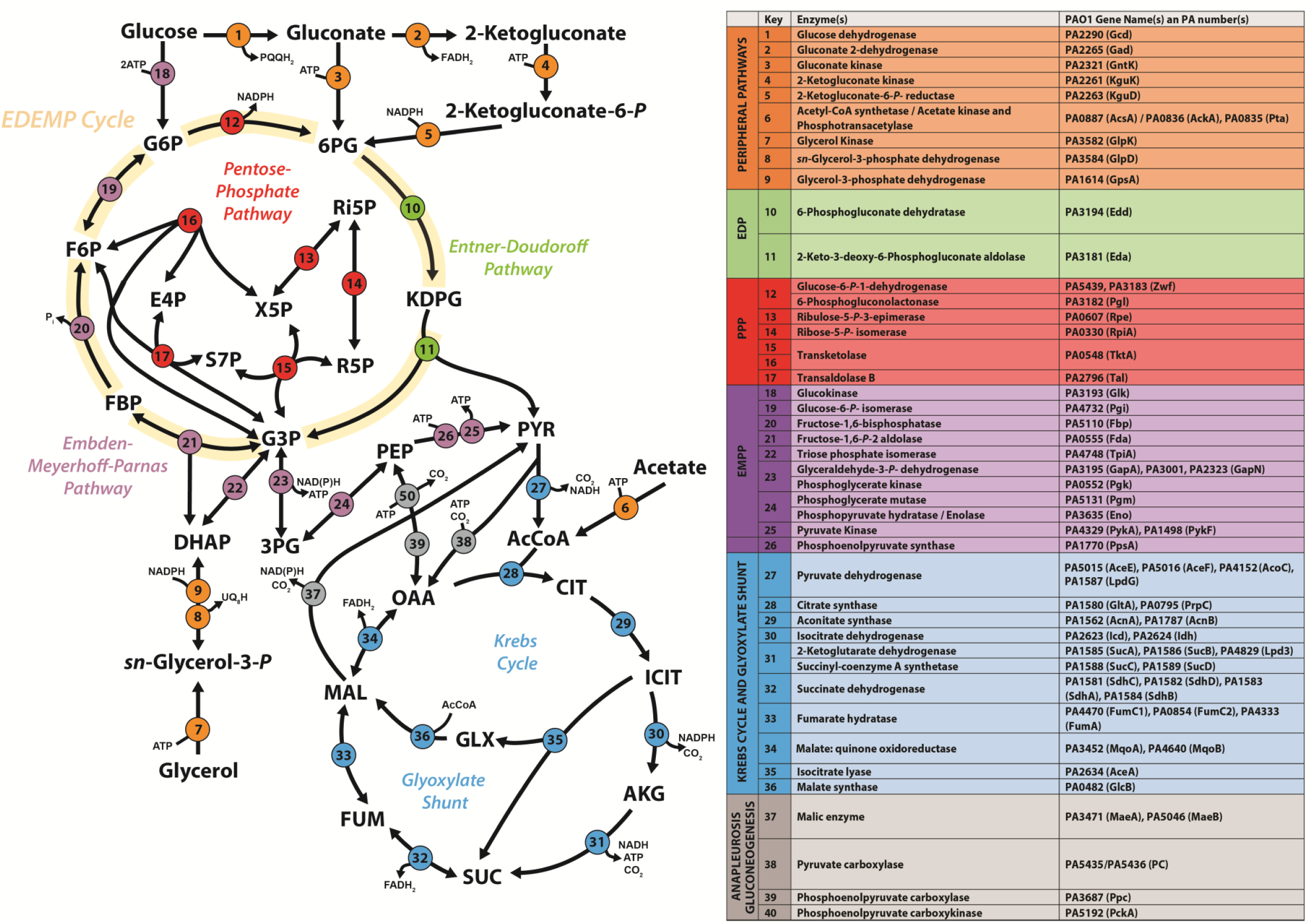
Biochemical pathways involved in central carbon catabolism in *P. aeruginosa* PAO1. The metabolic network was constructed around six main metabolic blocks, identified with different colours: (i) the peripheral pathways, that encompass the oxidative transformation of glucose, acetate and glycerol (orange); (ii) the Embden-Meyerhoff-Parnas pathway (EMP, non-functional in *P. aeruginosa* due to the absence of 6-phosphofructo-1-kinase, purple); (iii) the pentose phosphate pathway (PPP, red); (iv) the Entner-Doudoroff pathway (EDP, green); (v) the tricarboxylic acid cycle and glyoxylate shunt (blue); and (vi) anaplerotic and gluconeogenic bioreactions (grey). Figure adapted from [30].

RNA-Seq transcriptomic analyses on quadruplicate biological replicates yielded quantification of the mRNA levels from 5578 genes. After normalisation and statistical analysis (Figure S1 A-C), we identified 389 genes displaying increased expression on acetate, and 364 genes that displayed increased expression on glycerol (*p*-value ≤ 0.01, log_2_ fold change ≥2 or ≤-2) (File S1). A selection of these modulated genes were verified using promoter-luciferase transcriptional fusions (Figure S3) [14].

A complementary proteomic analysis reproducibly detected a total of 3921 proteins across three replicates for each condition. To our knowledge, this is the most comprehensive (in terms of coverage) transcriptome and proteome analysis carried out on *P. aeruginosa* to date. Following normalisation and statistical analysis (Figure S1), we identified 429 proteins that showed increased abundance during growth on acetate, and 402 proteins displaying increased expression on glycerol (*p*-value ≤ 0.01, log_2_ fold change ≥1 or ≤-1) (File S1). There was a clear relationship between the transcriptome and proteome fold-changes, particularly with respect to central carbon metabolism (Figure 2, File S3).

**Figure 2.**
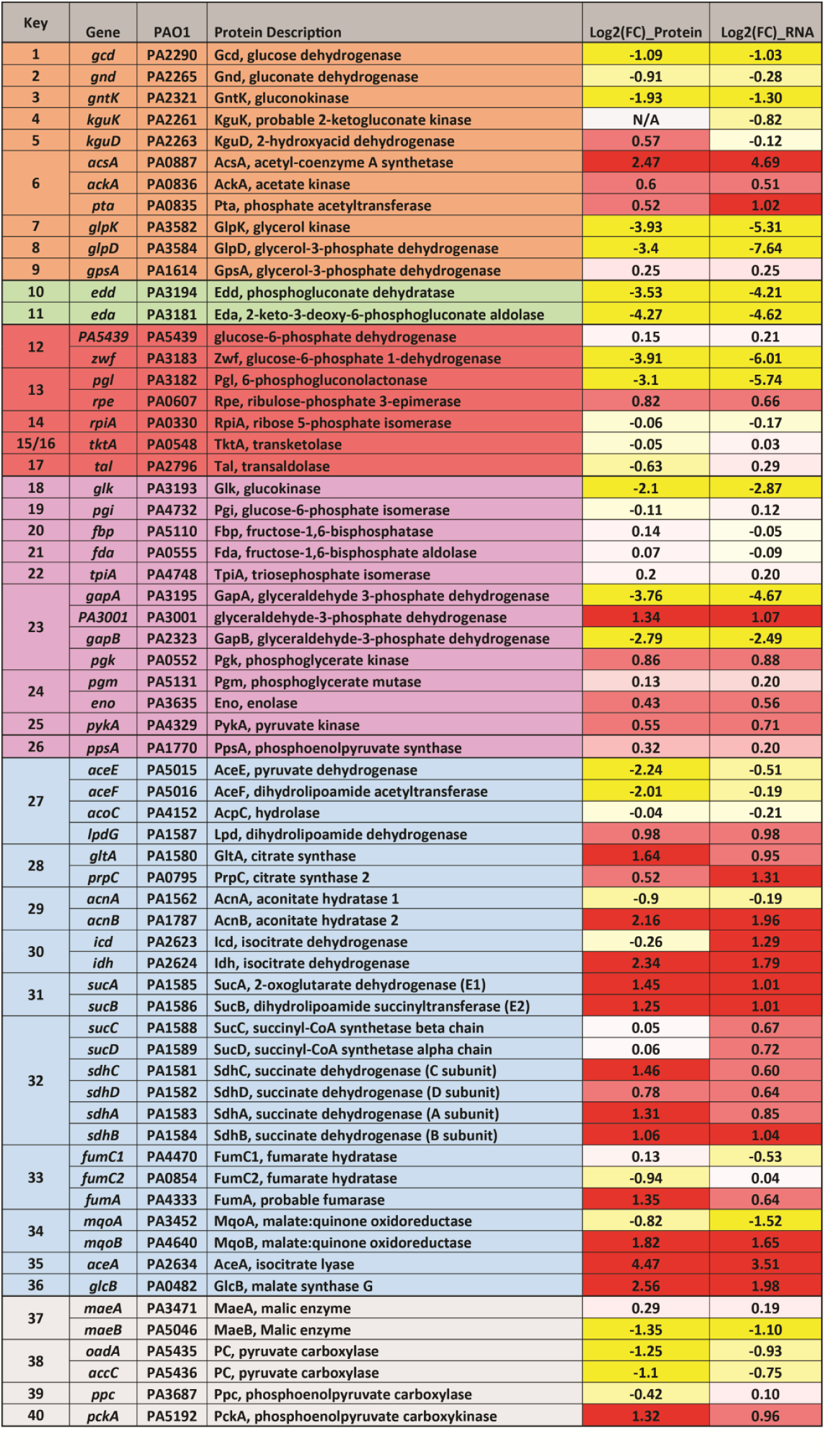
Comparison between protein and transcript fold-changes for selected *P. aeruginosa* enzymes involved in central carbon metabolism. The figure shows the log_2_ fold-changes in protein and transcript levels in (i) the peripheral pathways, that encompass the oxidative transformation of glucose, acetate and glycerol and the corresponding phosphorylated derivatives of these metabolites (orange); (ii) EMP pathway (non-functional, due to the absence of a 6-phosphofructo-1-kinase activity) (purple); (iii) the pentose phosphate (PP) pathway (red); (iv) the upper ED pathway (green); (v) the tricarboxylic acid cycle and glyoxylate shunt (blue); and (vi) anaplerotic and gluconeogenic bioreactions (grey). RNA-Seq and proteomic data are shown in File S1. Correlation plots are shown in File S3.

To allow for a more detailed analysis of the ‘omic alterations between these conditions, we used the proteomaps web service [15] to illustrate the statistically significant changes (*p*-value ≤ 0.01, log_2_ fold change ≥1 or ≤-1) as Voronoi tree maps. This provided a global overview of the proteomic consequences of growth in each carbon source. As shown in Figure 3 most of the proteomic changes were centred around ‘central carbon metabolism’, ‘biosynthesis’, ‘signalling and cellular process’ and ‘energy metabolism’.

**Figure 3.**
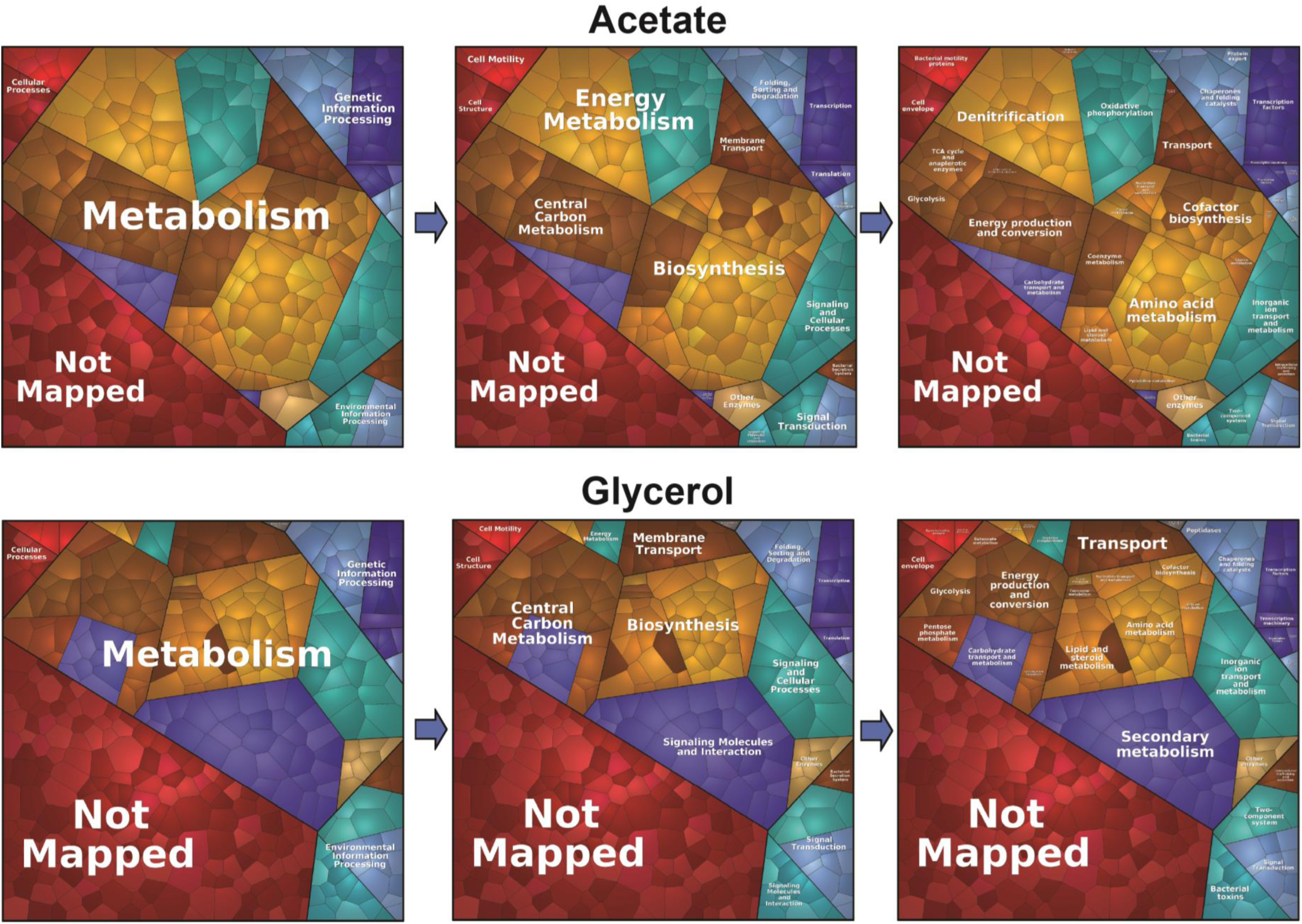
Growth on different carbon sources primarily affects metabolism. Illustration of the statistically significant changes (p-value ≤ 0.01, fold change ≥1 or ≤-1) during growth on glycerol and acetate as Voronoi tree maps using the proteomaps web service [15] Most of the proteomic changes were centred around ‘metabolism’, notably ‘central carbon metabolism’, ‘biosynthesis’, ‘signalling and cellular process’ and ‘energy metabolism’). Pathway assignment was performed using the Kyoto Encyclopedia of Genes and Genomes (KEGG) data set. Proteome alterations which could not be assigned to a specific pathway (uncharacterised/hypothetical proteins) are illustrated as ‘Not Mapped’.

Several recent studies have analysed the metabolic interactions of clinically-relevant bacteria within their host environment [16,17] and metabolic differences between mutant strains [18]. However, published fluxome studies for *Pseudomonas* species are scarce. In addition, all *P. aeruginosa* metabolic flux analysis (MFA) studies to date have been carried out using glucose as a sole carbon source [19–22], yet this substrate is not thought to play a major role during CF airway colonization [6,23,24]. To provide insight into the absolute metabolic fluxes of *P. aeruginosa* during growth on acetate and glycerol, we carried out a ^13^C fluxome analysis. This was done by measuring the mass isotopomer distributions of proteinogenic amino acids and cell carbohydrates (glycogen, glucosamine) using three separate tracers per carbon source (*Materials and Methods*) [25]. The calculated relative fluxes for the wild-type cultured on labelled glycerol or acetate are shown in Figure 4. Comparison of the flux maps, in combination with the proteomic/transcriptomic data, generated an unparalleled insight into the central carbon metabolic network(s) of *P. aeruginosa*.

**Figure 4.**
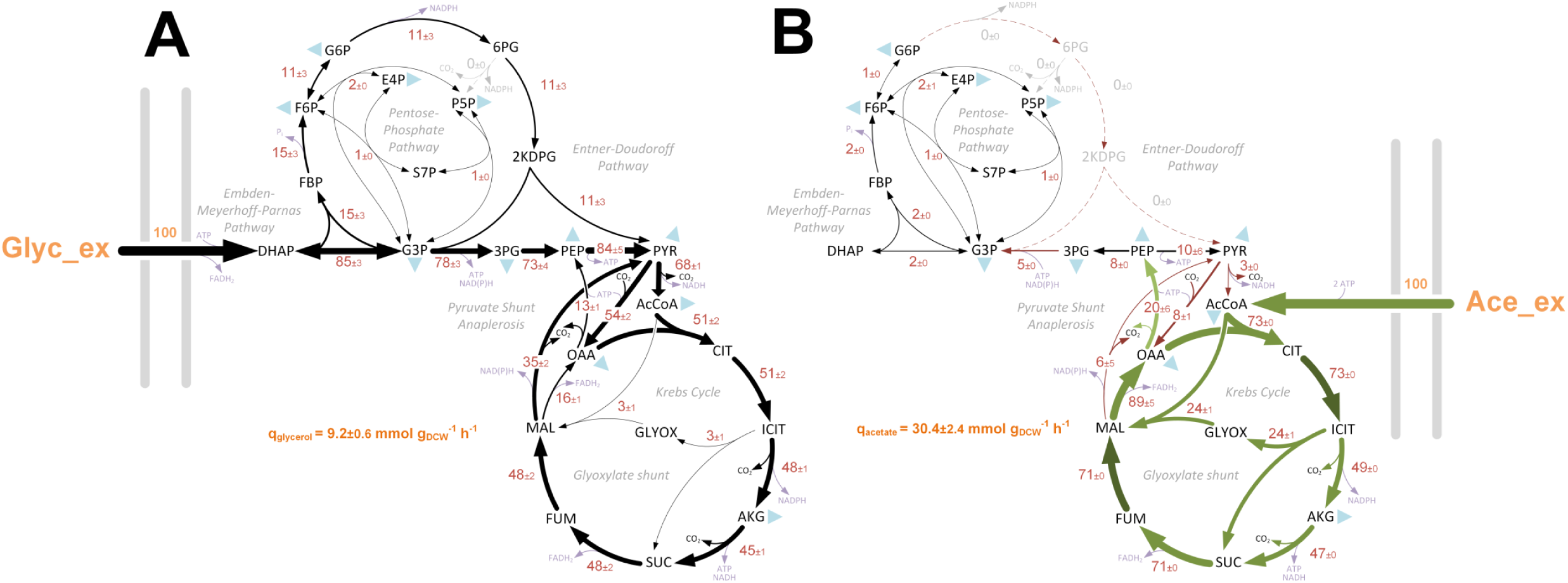
*In vivo* carbon flux distributions in central metabolism of *P. aeruginosa* PAO1 during growth on glycerol (A) or acetate (B) as sole carbon sources. Flux is expressed as a molar percentage of the average glycerol (9.2 mmol g^−1^ h^−1^) or acetate (30.4 mmol g^−1^ h^−1^) uptake rate, calculated from the individual rates in File S2. Anabolic pathways from 11 precursors to biomass are indicated as filled blue triangles. The flux distributions with bidirectional resolution (i.e., net and exchange fluxes), including the drain from metabolic intermediates to biomass and confidence intervals of the flux estimates, are provided in File S2. In agreement with previous studies of flux in *P. aeruginosa* and *Pseudomonas fluorescens*, we found no evidence for significant metabolite export during exponential growth in minimal media [19,21,22]. The errors given for each flux reflect the corresponding 90% confidence intervals. The full flux data sets are presented in Supporting Information File S2. Colours qualitatively indicate fluxomic correlation with changes on the protein/transcript level during growth in acetate (light green or red → significant up- or down-regulation (respectively); dark green or red → less significant up- or down-regulation).

#### Glycerol metabolism

During growth on glycerol, there was strong induction of the glycerol uptake system at both the proteomic and transcriptomic level (Figure 2, File S1). Glycerol is assimilated through the process of uptake (GlpF), phosphorylation (GlpK) and dehydrogenation (GlpD) to yield dihydroxyacetone phosphate (DHAP) [26–28]. Commensurate with this, the expression of all three enzymes/transporters (and their corresponding transcripts) was strongly stimulated during growth on glycerol.

Once synthesised, DHAP has two possible fates. One is anabolic. Here, DHAP is isomerised to generate glyceraldehyde 3-phosphate (G3P) through the action of triose phosphate isomerase (TpiA). DHAP and G3P are subsequently converted into fructose 1,6-*bis*phosphate (F1,6BP), thence to fructose 6-phosphate (F6P) and finally, to glucose 6-phosphate (G6P) through the action of Fda, Fbp and Pgi, respectively. This route is a necessary prerequisite for the generation of hexose sugars for cell wall synthesis and biomass production. The alternative fate of DHAP is catabolic. Here, G3P is converted to 1,3-*bis*phosphoglycerate for glycolytic energy production [29]. However, recent analyses of carbon fluxes in *Pseudomonas putida* suggests that the seemingly distinct anabolic and catabolic fates of DHAP/G3P may be more closely intertwined than previously thought, and that a proportion of the triose phosphate carbon skeletons are recycled rather than continuing to pyruvate *via* the lower glycolytic reactions [30]. This “EDEMP cycle” incorporates elements of the Entner-Doudoroff (ED) pathway, Embden-Meyerhof-Parnas (EMP) pathway, and Pentose Phosphate (PP) pathway. In glucose-grown *P. aeruginosa* and *P. putida* the EDEMP cycle operates in a markedly asymmetrical manner, with most (ca. 90%) of the flux proceeding through the ED pathway catalysed reactions; glucose → gluconate → 6-phosphogluconate → 2-keto-3-deoxy-6-phosphogluconate → G3P/pyruvate [25]. Intriguingly, and in spite of the anticipated demand for hexose synthesis, our data indicate that during growth on glycerol this asymmetry is retained, since the enzymes catalysing the “catabolic” reactions (Zwf, Pgl, Edd and Eda) are strongly up-regulated, whereas the enzymes catalysing the “anabolic” reactions (TpiA, Fda, Fbp and Pgi) remain relatively unaffected. We also note that glucokinase (Glk) and gluconokinase (GntK) were up-regulated during growth on glycerol. Intriguingly, this indicates that even in the absence of glucose, glycerol stimulates expression of the full ED glycolytic pathway [31].

Consistent with the proteomic/transcriptomic data, when *P. aeruginosa* was grown on glycerol, the fluxomics indicated that about 15% of the triose phosphates (G3P + DHAP) were diverted into the EDEMP cycle to form hexose phosphates [30]. Hexose generation is necessary for the synthesis of fructose 6-phosphate (F6P) and glucose 6-phosphate (G6P), required for biomass production. This recycling may also have a secondary function, since it generates NADPH through the action of glucose 6-phosphate dehydrogenase (Zwf). NADPH is a source of reducing agent for anabolism, and for dealing with oxidative insult [30]. However, the majority of the glycerol-derived carbon was oxidized to pyruvate through reactions in the lower half of the ED pathway. This notwithstanding, a significant amount of pyruvate was also generated from TCA-derived malate *via* the action of malic enzyme, feeding the so-called “pyruvate shunt”. The pyruvate shunt (a cyclical series of reactions converting malate → pyruvate → oxaloacetate → malate) generates NADPH for anabolism at the cost of consuming one equivalent of ATP. *P. aeruginosa* lacks phosphogluconate dehydrogenase activity (a major source of NADPH in many other species), so this apparently futile cycling reaction may be of major importance for growth. Indeed, > ⅔ of the carbon in the malate pool was shuttled back to pyruvate through the action of malic enzyme, with < ⅓ being converted to oxaloacetate *via* the malate dehydrogenase reaction. Overall, glycerol metabolism seems to be characterised by operation of the EDEMP cycle, net catabolism of G3P to pyruvate, and cycling of the latter through the pyruvate shunt.

#### Acetate metabolism

Growth on acetate as a sole carbon source elicited a similarly informative set of changes. There was a strong induction of the glyoxylate shunt enzymes (AceA and GlcB), which are known to be essential for growth on acetate [32,33], and of enzymes directly involved in acetate activation (such as AcsA, AckA and Pta) and acetate uptake (PA3234) [34]. The TCA cycle-associated enzymes were almost all up-regulated during growth on acetate, as was the membrane-bound malate-quinone oxidoreductase, MqoB (Figure 2). Unlike many bacteria, *P. aeruginosa* does not encode a soluble NADH-producing malate dehydrogenase, and MqoB directly donates the abstracted electrons to the membrane quinone pool [35].

Carbon fluxes in acetate-grown cultures were very different from those observed in glycerol-grown cells. Firstly, net flux through the lower reactions of the ED pathway was in the gluconeogenic direction. Second, the flow of carbon through the EDEMP cycle was much lower than that observed in glycerol, and flux terminated at the gluconeogenic end-point, G6P. The absence of flux through the G6P dehydrogenase (Zwf) reaction following this point is significant, since this reaction is often thought of as a major source of NADPH for anabolism. Third, and compounding this, the extent of carbon cycling through the NADPH-producing pyruvate shunt was low. Fourth, most of the carbon for anabolism was derived from oxaloacetate rather than malate. Presumably, this may reflect the increased rate of conversion of malate to oxaloacetate by the acetate-induced malate:quinone oxidoreductase (MqoB). However, perhaps the most notable difference was that during steady-state growth on acetate, around ⅓ of the carbon reaching the TCA cycle-glyoxylate shunt branchpoint was redirected into the glyoxylate shunt. [In glycerol-grown cells, only around 3% of the carbon reaching this branchpoint was redirected to the glyoxylate shunt.] The glyoxylate shunt serves to supply the cell with malate and thence (following the MqoB-catalyzed conversion) also the gluconeogenic precursor, oxaloacetate. In an elegant feedback loop, high levels of oxaloacetate (such as would accumulate if the NADPH supply for anabolism was limiting) stimulate the activity of one of the isocitrate dehydrogenase isozymes, IDH, thereby restoring flux through the TCA cycle [36]. One outcome of this is that NADPH levels become replenished because the IDH-catalyzed reaction is a major source of this coenzyme *in vivo*. This presumably relieves the limitation on oxaloacetate usage in anabolism. At the steady state, our flux data show that around ⅔ of the carbon flowing through the branchpoint is fluxed through the isocitrate dehydrogenases.

Our flux data show that no carbon passes through the NADPH-generating steps of the ED pathway during growth on acetate. So far, we have primarily framed our assessment of these observations around the need for NADPH in anabolism. However, it has not escaped our attention that NADPH limitation may impact on infection too. Several earlier workers have suggested that flux through the ED pathway may confer an additional benefit on *P. aeruginosa* by providing sufficient reducing power (NADPH) to counteract host-mediated oxidative stress [37,38]. Whereas our data neither confirm nor refute this notion, we show here that four major NADPH-supplying reactions are accessible, depending on the substrate; transhydrogenation (NADH → NADPH), the EDEMP cycle, the pyruvate shunt (malic enzyme), and the isocitrate dehydrogenase(s)-catalysed reaction.

### Beyond central metabolism – growth in acetate induces extensive remodelling of the electron transport chain

Aerobic respiration re-oxidizes NADH, thereby generating energy, maintaining redox homeostasis, and ensuring continued oxidative metabolism [39] [40]. Bacteria are known to coordinate the composition of the electron transport chain (ETC), and in particular, levels of the terminal oxidases, according to their metabolic needs [41]. The ETC of *P. aeruginosa* contains three NADH dehydrogenases; the multi-subunit proton pumping NDH-1 (encoded by *nuoA-N* (PA2637-PA2649)), the HQNO-resistant proton pump Nqr (encoded by *nqrA-F* (PA2994-2999)), and a single-subunit flavoenzyme which does not participate in ion translocation (NDH-2, encoded by *ndh*, PA4538) [42]. All three dehydrogenases displayed increased expression during growth on acetate, particularly at the protein level (file S1).

A characteristic feature of the *P. aeruginosa* respiratory chain is its use of high-affinity terminal oxidases, even during aerobic growth. This unusual wiring of the *P. aeruginosa* ETC is thought to drive the formation of a microaerobic environment; a trait that may give the pathogen a fitness advantage during infection [43]. Expression of the terminal oxidases in *P. aeruginosa* is directly or indirectly controlled by a two-component system, RoxS-RoxR. The kinase, RoxS, is thought to sense the redox status of the respiratory chain, either by titrating the redox status of the ubiquinone/ubiquinol (UQ) pool or by responding to electron flux through the terminal oxidases. UQ acts as the electron donor for complex III, the quinol oxidases (Cyo), and the cyanide-insensitive terminal oxidase, CIO. Complex III transfers electrons to a c-type cytochrome, which then acts as the electron donor for the terminal cytochrome oxidases Cox, Cco1, or Cco2. Cco1 is constitutively expressed at high levels, whereas Cco2 is thought to support growth under low oxygen tension (∼2% O_2_), as it is regulated by the anaerobic sensor, Anr [44]. The ‘omics data indicated that Cco1, Cco2, Cyo and complex III subunits (encoded by PA4429-PA4431) are highly-expressed during growth on acetate, whereas the Cox oxidase was more highly-expressed during growth on glycerol. This may be a strategy to limit oxidative stress, as Cox has a high H^+^/e^−^ratio and can extract more energy per unit of carbon [45,46]. The greater expression of Cox during growth on glycerol may be a consequence of RoxSR-mediated inhibition of *cox* gene expression during growth on acetate (File S1) [47].

Levels of the soluble pyridine nucleotide transhydrogenase, Sth (PA2991), were increased ca. 3-fold during growth on acetate. Pyridine nucleotide transhydrogenases catalyse the reversible reduction of either NAD^+^ or NADP^+^ by NADPH or NADH (respectively). The primary physiological role of Sth is thought to be in the NAD^+^-dependent re-oxidation of NADPH [48]. Re-oxidation of excess NADPH is likely to be important during growth on acetate, as extensive catabolism of this substrate through the TCA cycle generates more NADPH than is required for biosynthesis [49]. Interestingly, the NAD(P) transhydrogenase encoded by *pntAA* and *pntAB* was not differentially expressed during growth on acetate or glycerol, suggesting that Sth is the main transhydrogenase used by *P. aeruginosa* in these conditions [48].

As noted earlier, *P. aeruginosa* cultured in glycerol has a significantly slower growth rate than cells grown in acetate, glucose or succinate (Figure S2). This lower growth rate may result in a lower metabolic demand, meaning these cells do not require high expression of terminal oxidases and other ETC components. By contrast, higher growth rates may result in accumulation of NADH due to the rate-limitations inherent in ETC-dependent re-oxidation. This is because faster growing cells are known to increase their length [53], suggesting that they are likely have lower surface-to-volume ratios. This limits the membrane’s physical capacity for accommodating respiratory complexes, and thereby also limits NADH re-oxidation [50]. To investigate this possibility further, we examined the length of exponentially-growing *P. aeruginos*a cells grown in MOPS media with various carbon sources. To do this, we introduced an eGFP-expressing plasmid (pMF230) to PAO1 and cultured the strain in MOPS medium containing different carbon sources [51]. When cells reached an OD_600_ of 0.5, they were fixed and examined by fluorescence microscopy. As shown in Figure S6 (and File 3), the slower-growing cells in MOPS-glycerol were indeed significantly shorter than cells grown in acetate, glucose or succinate [52]. This supports the notion that *P. aeruginosa* cell size (and hence, surface-to-volume ratio) is dependent on growth rate.

### Induction of the denitrification apparatus in *P. aeruginosa* during aerobic growth

When O_2_ is limiting, *P. aeruginosa* can use nitrate as an alternative electron acceptor. This is made possible by the presence of nitrate reductase (NAR) which can accept electrons from the UQ pool, and nitrite reductase (NIR), which receives electrons *via* complex III and cytochrome c [53]. Denitrification is thought to be an important pathway to support the anoxic growth of *P. aeruginosa* in the CF airways [54,55] and nitrate is a known component of human body fluids, arising from diet and NO auto-oxidation. Indeed, NO_3_ ^−^ concentrations in the CF airways have been reported between 73 μM – 792 μM, with an average of 348 μM [56]. Molecular oxygen is therefore not essential for growth in this environment.

The gene clusters encoding the dissimilatory nitrate reductases (*nar* and *nap*) and those encoding nitrous oxide (N_2_O) utilisation (*nos*) are unlinked, whereas the genes encoding nitrite respiration (*nir*) and nitric oxide respiration (*nor*) are adjacent. In addition to the enzymes above, the denitrification operons also harbour genes for ancillary functions such as cofactor synthesis, transport and protein maturation [53]. Both the proteomic and transcriptomic analyses showed activation of the denitrification pathways following growth in MOPS-acetate (File S1). This was surprising, since the cultures were harvested during exponential growth in baffled, well-aerated conical flasks without added nitrate. This is an important point, because denitrification in *P. aeruginosa* is known to be under the control of the master regulator Dnr. In turn, Dnr is under the control of the anaerobic transcriptional regulator Anr. The latter contains an oxygen sensitive [4Fe-4S]^2+^ cluster and is thought to be active only under conditions of oxygen limitation. In contrast, Dnr is a nitric oxide (NO) sensor, and contains a ferrous heme. Dnr protein expression was up-regulated 2.8-fold during growth on acetate compared with glycerol (File S1). Previous work has shown that, in addition to Dnr-mediated regulation, the *narGHJI* operon is also activated by the NarXL two-component system. The NarX and NarL proteins were up-regulated 2-fold during growth on acetate. Taken together, these data indicate that in spite of the presence of molecular oxygen and the absence of added nitrate, growth on acetate strongly induces the denitrification apparatus in comparison with glycerol. We conclude that the denitrification apparatus and components required for aerobic respiration are expressed simultaneously during growth on acetate [57].

### *P. aeruginosa* cellular NAD(P)(H) ratios change in a carbon source dependent manner

By summing the fluxes through reactions that either produce or consume NADPH or ATP, we were able to calculate the relative contributions of different metabolic reactions to the redox (NADPH) and energy (ATP) balances during growth on each carbon source (Figure 5). This revealed that cells grown on glycerol obtain their NADPH from a variety of reactions, whereas acetate-grown cells are heavily-reliant on NADPH derived from the isocitrate dehydrogenase(s) reaction. Moreover, most of the NADPH generated during growth on acetate was used in anabolism. With regards to ATP, cells grown on glycerol were effectively “over-supplied” with this reagent, maintaining a substantial “operating surplus” of ATP. In contrast, the supply of ATP in cells grown on acetate was essentially identical to the metabolic demand. The most likely reason for this is the 2 x ATP ‘cost’ of acetate uptake and activation [58]. During growth on glycerol, the cell generates 2.6 moles of ATP per mole of C, whereas just 1.1 moles of ATP are generated on acetate. This “energy deficit” may explain why the cell induces additional mechanisms to maximise energy production during growth on acetate (e.g., high affinity electron acceptors, increased denitrification, increased Sth transhydrogenase expression etc (File S1)).

**Figure 5.**
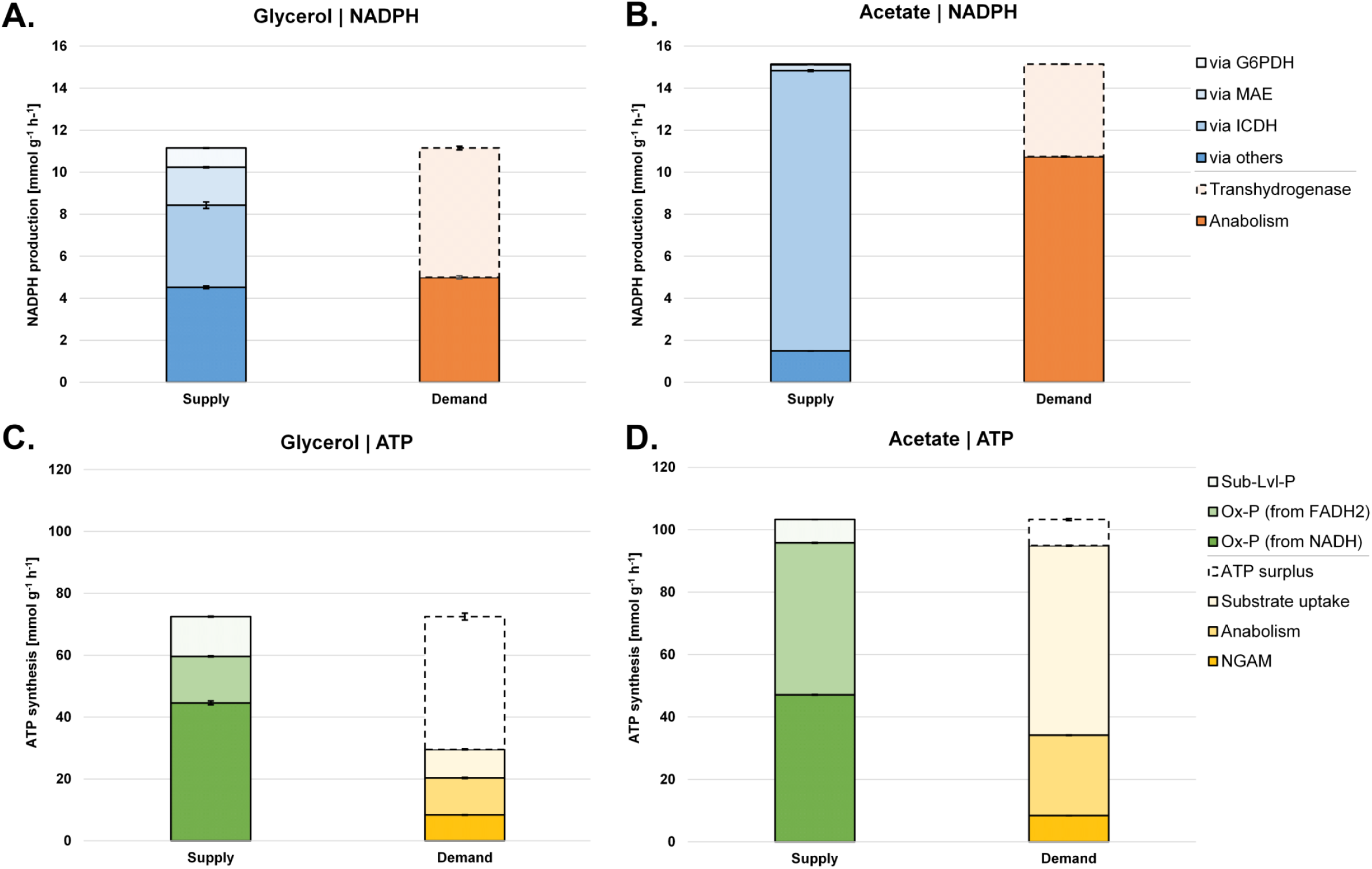
Quantitative analysis of NADPH supply and demand (redox) for glycerol (A) and acetate (B) grown *P. aeruginosa*. ATP (energy metabolism) supply and demand for glycerol (A) and acetate (B) grown *P. aeruginosa*. 6). Reactions linked to NADPH (A-B) and ATP (C-D) metabolism were calculated from the obtained fluxes (Fig. 4). They are given as absolute fluxes (mmol g^−1^ h^−1^) and are related to the specific carbon uptake rate (see File S2). G6PDH; glucose 6-phosphate dehydrogenase, MAE; malic enzyme, ICDH; isocitrate dehydrogenase, Ox-P; oxidative phosphorylation, NGAM; non-growth associated maintenance needs.

We anticipated that the faster growth rate (Figure S2) and TCA cycle driven metabolism (Figure 2) during growth on acetate would result in an increased rate of respiration, and that this might drive the NAD(P)H/NAD(P)^+^ ratio towards a more reduced state [50]. This was indeed the case (Figure S2) indicating that carbon source alone is sufficient to alter the intracellular redox economy of *P. aeruginosa*. We next investigated what might be driving this increased NADH/NAD^+^ ratio during growth on acetate. Microorganisms display an elevated NADH/NAD^+^ ratio when NADH re-oxidation is slowed by limitation of electron acceptors [59], or when the metabolism of certain carbon sources outpaces the capacity of the *P. aeruginosa* ETC to re-oxidise the coenzyme. This may explain why cells grown in acetate appear to scavenge for alterative electron acceptors (such as nitrate, as indicated by the increased expression of the denitrification apparatus). To determine if aerobic denitrification might be used by *P. aeruginosa* to correct the reduced status of its redox pool, we therefore examined the impact of nitrate addition on the NADH/NAD^+^ ratio. As shown in Figure 7, the addition of 20 mM nitrate to MOPS-acetate cultures decreased the NADH/NAD^+^ ratio during exponential growth. Absolute quantitation of the NADH and NAD^+^ levels (Figure S2) confirmed that the total NAD(H) and NADP(H) pools were similar in the presence and absence of added nitrate. To confirm that this drop in the NADH/NAD^+^ ratio was due to aerobic denitrification, we generated a deletion mutant in the master transcriptional regulator of denitrification, *dnr*. As expected, nitrate reductase (NirS) expression was abolished in the *dnr* mutant, and also in an *anr* mutant (Anr controls the expression of *dnr*) (Figure S4). Crucially, and consistent with findings from other laboratories showing that nitrate addition has little impact on the NADH:NAD^+^ ratio in mutants defective in denitrification [60], the *dnr* mutant was also defective in nitrate-dependent re-oxidation of NADH and NADPH (Figure 7). We conclude that aerobic denitrification during growth on acetate impacts on the redox status of the NAD(P)H/NAD(P)^+^ pool.

**Figure 6.**
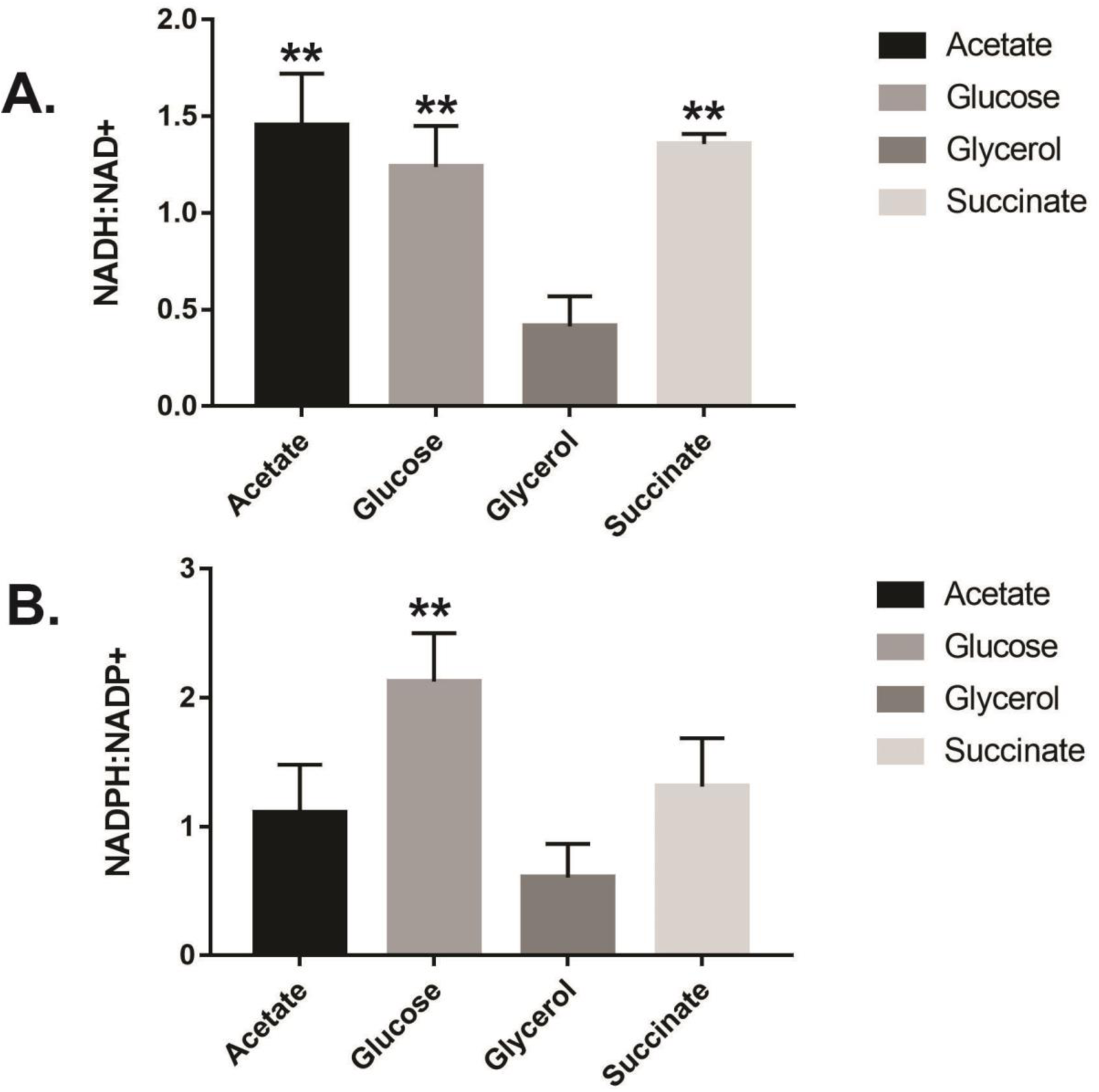
Maximal measured NADH:NAD^+^ and NADPH:NADP^+^ ratios in *P. aeruginosa* grown in the indicated sole carbon sources. Exponentially-growing cells in MOPS-glycerol have a significantly lower NADH:NAD^+^ ratio (p < 0.01) compared with cells grown in MOPS-acetate, MOPS-glucose or MOPS-succinate, and a significantly lower NADPH:NADP^+^ ratio compared with cells grown in MOPS-glucose (p = 0.0064). Total NAD(P)(H) concentrations at each time-point and in each carbon source are shown in Figures S2 and S4. The data were analysed using GraphPad Prism (v 6.01) using t-test statistical analysis (MOPS-glycerol *versus* MOPS-acetate, MOPS-glucose or MOPS-succinate).

**Figure 7.**
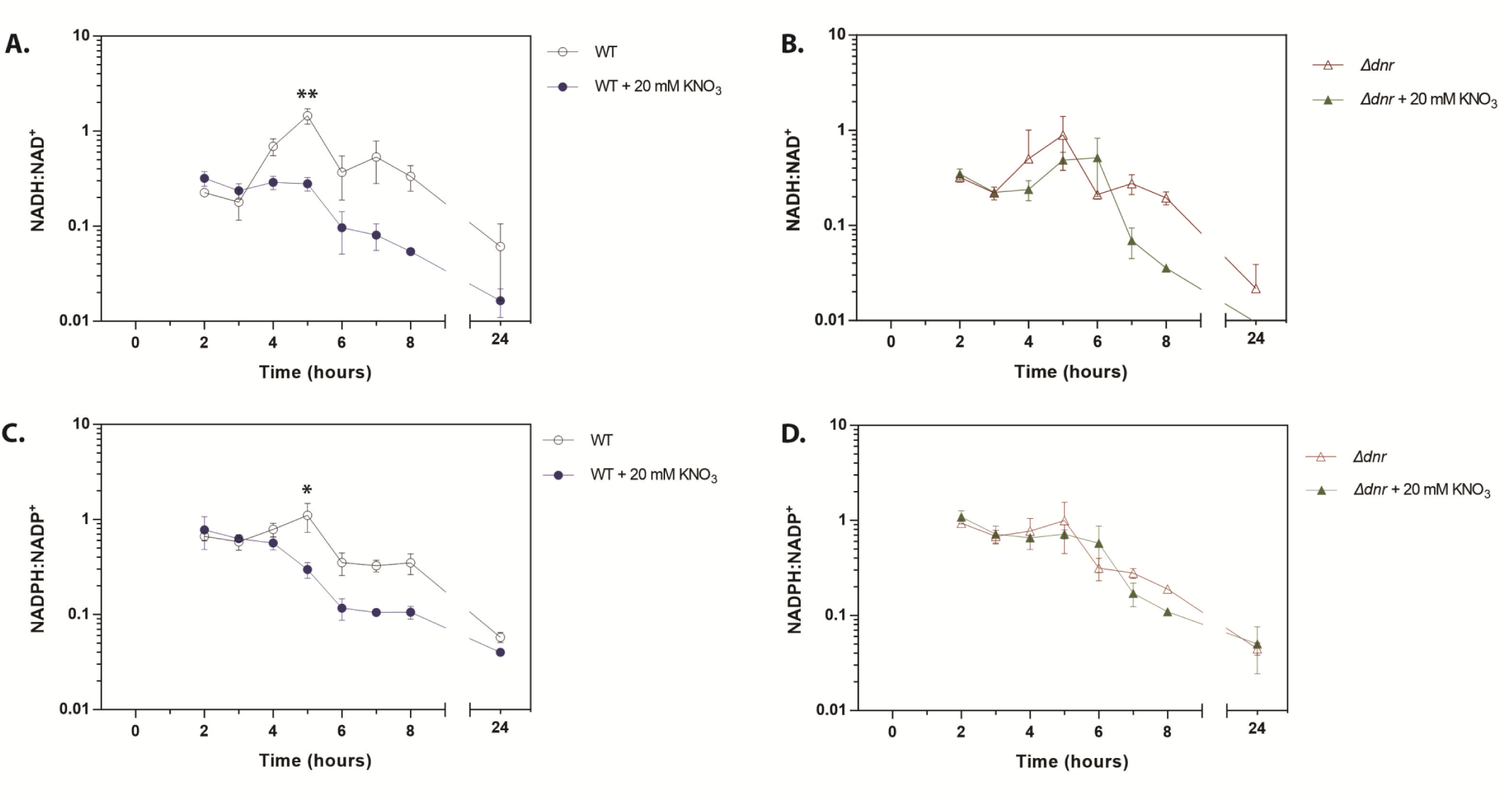
Coenzyme re-oxidation is impaired in a *dnr* mutant. The NADH:NAD^+^ (A and B) and NADPH:NADP^+^ (C and D) ratios were measured in cultures of wild-type PAO1 (A and C) and in cultures of an isogenic Δ*dnr* mutant (B and D). Cultures were grown in MOPS-acetate -/+ 20 mM KNO_3_, as indicated. Corresponding CFUs are shown in Figure S4. The data were analysed using GraphPad Prism (v 6.01) using t-test statistical analysis at 5h growth. Nitrate addition to PAO1 wild-type resulted in a significant reduction in the NADH:NAD^+^ ratio (p = 0.0017, indicated by **) and the NADPH:NADP^+^ ratio (p = 0.0202, indicated by *). Nitrate addition to a Δ*dnr* mutant did not significantly alter either ratio (p = 0.8065, p = 0.1862).

## Discussion

Several earlier studies have described the systems-level cellular adjustments that accompany the growth of industrially-relevant model organisms such as *E. coli, Saccharomyces cerevisiae, B. subtilis, C. glutamicum, B. succiniciproducens*, and *P. putida* in single carbon sources [61–66]. However, and although the accrued metabolic models have made a valuable contribution towards our “textbook understanding” of microbial metabolism, it is clear that they can be only loosely extrapolated to other organisms, such as *P. aeruginosa* [67]. Surprisingly, a comparative ‘omics study has yet to be carried out for *P. aeruginosa* grown on carbon sources relevant to infection. Overall, carbon preference remains poorly understood for *P. aeruginosa*, even though it is known to have a profound impact on virulence-associated phenotypes, including toxin production, biofilm formation and growth rate. In this work, we rectify this by developing a high-resolution global map of *P. aeruginosa* metabolism during growth on two infection-relevant carbon sources, acetate and glycerol.

Our data indicate a clear relationship between transcript and protein changes in *P. aeruginosa* for the growth regimens tested [20,36,68]. These expression data enabled us to establish which metabolic pathways are active in each growth condition, thereby providing an experimentally-supported framework for interpreting the fluxomic data. This notwithstanding, the fluxomics highlighted the possibility of additional layers of regulatory complexity involved in fine-tuning of central carbon metabolism. For example, growth on acetate results in the expression of three enzymes (the isocitrate dehydrogenases, ICD and IDH, and the isocitrate lyase, ICL), all of which compete for a shared substrate, isocitrate. These enzymes are known to exhibit radically different catalytic and regulatory properties compared with their *E. coli* counterparts [36]. In spite of this, our fluxomic analysis revealed a remarkably similar flux partitioning at the branchpoint between the TCA cycle and the glyoxylate shunt in both organisms [36].

In addition to alterations in central carbon metabolism, we also noted substantial remodelling of the ETC composition during growth in acetate compared with glycerol. For example, we noted a substantial increase in the expression of terminal oxidases (*cyo, cco1* and *cco2*) during growth on acetate, and increased *cox* expression during growth on glycerol. This suggests that different terminal oxidases are employed in different growth conditions, allowing metabolism to be optimally adjusted for energy generation [69]. Interestingly, growth on mucus has been previously shown to stimulate Cyo expression and to repress Cco2 expression [70]. Our data suggest that these observations may be driven by metabolism of one or more mucus-derived compounds.

The ETC alterations that we observed included strong aerobic induction of the denitrification machinery during growth on acetate. This was unexpected, since denitrification is usually associated with oxygen limitation and microaerobic growth. The signal which leads to this remodelling of the ETC remains to be elucidated. However, we speculate that these changes in ETC architecture and composition may be a mechanism to restore redox balance (NADH:NAD^+^ ratio) in the cell [71]. Acetate-grown cells accumulate NADH, so activating alternative mechanisms (such as the denitrification apparatus) to oxidise NADH may be a homeostatic attempt to regain an optimal cellular redox status. Consistent with this, the addition of nitrate to acetate-grown *P. aeruginosa* did lead to a more oxidized NADH:NAD^+^ ratio. Moreover, a mutant (Δ*dnr*) defective in denitrification, was unable to maintain optimal redox homeostasis. However, growth was not affected in the Δ*dnr* mutant, perhaps suggesting that the organism also has additional (possibly compensatory) mechanisms to deal with redox dysregulation [37,72].

Aerobic denitrification has been recently suggested to function as a ‘bet-hedging’ strategy to anticipate nitrate availability and respond to abrupt anoxia [73,74]. The oxygen-limited microenvironment in the CF airways is thought to result in *P. aeruginosa* growth by microaerobic respiration [75]. One possibility is that hybrid respiration (a combination of aerobic respiration and denitrification) may well be a predominant mechanism of survival in these conditions [76]. Genes under the control of the transcriptional regulator Anr are also abundant in *P. aeruginosa* RNA extracted from CF sputum [77–79].

A key fundamental question arises from this study – how does *P. aeruginosa* activate the denitrification system during growth in aerobic conditions? Until now, induction of the denitrification machinery has been largely attributed to the activity of Anr (and its subordinate denitrification-specific regulator, Dnr). Anr dimerization is thought to be dependent on the formation of an oxygen-liable [4Fe-4S]^2+^, which is destabilised and disassociates in the presence of oxygen, abolishing Anr activity. However, there is accruing evidence that Anr may be active even in the presence of oxygen. For example, Jackson *et al*., have shown that choline catabolism leads to aerobic expression of the Anr regulon, suggesting that Anr activity can be sustained in the presence of oxygen. Also, Anr overexpression in well-aerated cultures increases the levels of *dnr* transcript 29-fold [80]. Furthermore, the expression of *nir* and *nar* genes has been shown to increase in late exponential phase cultures compared with stationary phase cultures of PA14 [81]. This mobilisation of the Anr regulon in the presence of oxygen suggests that Anr can modulate gene expression without fully intact metallo-centres or dimers. This may be to assist the cells during the transition to conditions of low oxygen tension [80].

Finally, our data demonstrate that *P. aeruginosa* grows more rapidly on acetate than it does in glycerol, both in terms of specific growth rate (measured as optical density) and cell length (Figure S6). In order to maintain redox homeostasis in such conditions, cells will often switch to “overflow metabolism” whereby partially oxidised metabolic intermediates such as acetate are excreted [59,82]. At first glance, overflow metabolism seems wasteful, although it does allow them to maximise their growth rate [83]. It is tempting to speculate that aerobic denitrification in *P. aeruginosa* may also be a form of ‘overflow’ metabolism in which the alternate electron acceptor nitrate is utilised alongside oxygen in an effort to maintain redox homeostasis.

## Materials and Methods

### Growth Conditions

Unless otherwise indicated, *P. aeruginosa* strain PAO1 [84] was routinely grown in lysogeny broth (LB LENNOX) (Oxoid Ltd) at 37°C with shaking (250 rpm). The strains used in this study are listed in Table S1. The overnight pre-cultures were started from separate clonal source colonies on streaked LB agar plates. Strains were cultured in MOPS (morpholinepropanesulfonic acid) media with the relevant carbon sources [85]. Cell growth was monitored as optical density in a spectrophotometer at a wavelength of 600 nm (OD_600_). A previously determined conversion factor of 0.42 g CDW per OD unit was used to calculate biomass specific rates and yields from the obtained OD_600_ values [21].

### Transcriptomics (RNA-Seq)

PAO1 was grown in 40 mL MOPS with acetate or glycerol as sole carbon sources (quadruplicate), 37°C with shaking (250 rpm) in baffled flasks (500 mL). An aliquot (5 mL) of culture was harvested from each sample at OD_600_ of 0.5 (exponential growth) and added to an equal volume of RNAlater®. Samples were then sedimented in an Eppendorf 5810R centrifuge at 3220 × *g* for 15 min (4°C) and the pellets were stored at –80°C. Total RNA was isolated as described in [86], followed by phenol-chloroform-isoamyl alcohol extraction and ethanol precipitation. Ribosomal RNA (rRNA) was then depleted from each sample (5 µg each) using the bacterial Ribo-Zero rRNA Removal Kit (Illumina). The integrity of the RNA was evaluated using an RNA 6000 Nano LabChip and an Agilent 2100 Bioanalyzer (Agilent Technologies, Germany). Eight indexed, strand-specific cDNA libraries were prepared, and samples were sequenced on an Illumina HiSeq 2000 with a 51 bp single-end read length (GATC Biotech, Germany). The sequencing data are deposited at ArrayExpress (accession number E-MTAB-8374).

### Reads mapping and annotations

The resulting FASTQ files were mapped to the PAO1 genome obtained from the Pseudomonas Genome Database (PDG) (http://www.pseudomonas.com/) using TopHat v.2.0.3 [37], Bowtie v.0.12.8 [38] with a ∼97% success rate to generate SAM files. The sequence reads were adaptor clipped and quality trimmed with trimmomatic [87] with default parameters. Transcript abundance and differential gene expression were calculated with the program Cufflinks v.2.0.1 [88]. Gene expression levels were normalized using fragments per kilobase of exon per million mapped reads (FPKM) report values. Genes were considered as induced or repressed, only when their log_2_ fold change was >1 or <−1, respectively, and their *p*-value was <0.01 (File S1, Figure S1).

### Quantitative proteomic analysis

*P. aeruginosa* cells (OD_600_ = 0.5, 30 mL) were harvested in identical growth conditions as the transcriptomics section described above. Pellets were resuspended in lysis buffer (100 mM Tris-HCl, 50 mM NaCl, 10% (v/v) glycerol, 1 mM tris(2-carboxyethyl)phosphine (TCEP), pH 7.5) with cOmplete Mini protease inhibitor cocktail (Roche). Following three rounds of sonication (3 × 10 sec) on ice, supernatants were clarified by sedimentation (21130 × *g*, 15 min, 4°C) in an Eppendorf 5424R centrifuge. Aliquots (100 μg) of each sample was reduced with TCEP, alkylated with iodoacetamide and labelled with Tandem Mass Tags (TMTs). TMT labelling was performed according to the manufacturer’s protocol (https://www.thermofisher.com/order/catalog/product/90110).

### LC-MS/MS

Dried fractions from the high pH reverse-phase separations were resuspended in 30 µL of 0.1% (v/v) formic acid (14 combined fractions). LC-MS/MS experiments were performed using a Dionex Ultimate 3000 RSLC nanoUPLC (Thermo Fisher Scientific Inc, Waltham, MA, USA) system and a Lumos Orbitrap mass spectrometer (Thermo Fisher Scientific Inc, Waltham, MA, USA). Peptides were loaded onto a pre-column (Thermo Scientific PepMap 100 C18, 5 mm particle size, 100 Å pore size, 300 mm i.d. x 5 mm length) from the Ultimate 3000 auto-sampler with 0.1% (v/v) formic acid for 3 min at a flow rate of 10 µL/min. Separation of peptides was performed by C18 reverse-phase chromatography at a flow rate of 300 nL/min using a Thermo Scientific reverse-phase nano Easy-spray column (Thermo Scientific PepMap C18, 2 mm particle size, 100 Å pore size, 75 mm i.d. x 50 cm length). Solvent A was water + 0.1% formic acid and solvent B was 80% (v/v) acetonitrile, 20% water + 0.1% formic acid. The linear gradient employed was 2-40% B in 93 min (total LC run time was 120 min including a high organic wash step and column re-equilibration).

The eluted peptides were sprayed into the mass spectrometer by means of an Easy-Spray source (Thermo Fisher Scientific Inc.). All *m/z* values of eluting peptide ions were measured in an Orbitrap mass analyser, set at a resolution of 120,000 and were scanned between *m/z* 380-1500 Da. Data dependent MS/MS scans (Top Speed) were employed to automatically isolate and fragment precursor ions by collision-induced dissociation (CID, Normalised Collision Energy (NCE): 35%) which were analysed in the linear ion trap. Singly charged ions and ions with unassigned charge states were excluded from being selected for MS/MS and a dynamic exclusion window of 70 sec was employed. The top 10 most abundant fragment ions from each MS/MS event were then selected for a further stage of fragmentation by Synchronous Precursor Selection (SPS) MS3 [89] in the HCD high energy collision cell using HCD (High energy Collisional Dissociation, (NCE: 65%). The *m/z* values and relative abundances of each reporter ion and all fragments (mass range from 100-500 Da) in each MS3 step were measured in the Orbitrap analyser, which was set at a resolution of 60,000. This was performed in cycles of 10 MS3 events before the Lumos instrument reverted to scanning the *m/z* ratios of the intact peptide ions and the cycle continued.

### Proteomic Data Analysis

Proteome Discoverer v2.1 (Thermo Fisher Scientific) and Mascot (Matrix Science) v2.6 were used to process raw data files. Data was aligned with the UniProt *Pseudomonas aeruginosa* (5584 sequences) the common repository of adventitious proteins (cRAP) v1.0.

The R package MSnbase [90] was used for processing proteomics data. Protein differential abundance was evaluated using the Limma package [91]. Differences in protein abundances were statistically determined using the Student’s *t*-test with variances moderated by Limma’s empirical Bayes method. *P*-values were adjusted for multiple testing by the Benjamini Hochberg method [92]. Proteins were considered as increased or decreased in abundance, only when their log_2_ fold change was >1 or <−1, respectively, and their *p*-value was <0.01. The mass spectrometry proteomics data have been deposited to the ProteomeXchange Consortium *via* the PRIDE (55) partner repository with the data set identifier PXD015615.

### Construction of Luciferase reporter strains

Translational reporter constructs were made by fusing the upstream promoter sequences with *luxCDABE* using the primers listed in Table S1. Purified PCR products were digested and directionally ligated into the multiple cloning site of the pUC18T-mini-Tn7T-lux-Gm plasmid [14]. The mini-Tn7-lux element were integrated into the PAO1 chromosome by electroporation along with the helper plasmid pTNS2 as previously described [93].

### Luciferase-Promoter Assay

Luciferase and OD_600_ readings were measured using a BMG Labtech FLUOstar Omega microplate reader. Strains were cultured in MOPS media with the relevant carbon sources (100 µL) in 96 well microplates (Greiner bio-one, F-Bottom, Black), covered with gas permeable imaging seals (4titude - 4ti-0516/96). Luciferase expression was assessed every 30 min (Gain = 3600) for up to 24 hr. Growth was assessed by taking OD readings at 600 nm simultaneously with the luminescence readings Luciferase readings were expressed as relative luminometer units (RLU) normalised to OD_600_ in order to control for growth rate differences across the selected carbon sources.

### ^13^C fluxomics

Starter cultures were grown In LB medium. For the second and main cultures, PAO1 was grown in MOPS minimal media with 60 mM acetate or 40 mM glycerol as the sole carbon source (120 mM carbon). For ^13^C flux experiments, naturally labelled acetate and glycerol was replaced with three separate tracers per carbon source to maximise dataset resolution and to accurately determine substrate uptake. Naturally labelled glycerol was substituted with [1,3-^13^C_2_] glycerol (99%), [2-^13^C] glycerol (99%) and an equimolar mixture of [U-^13^C_3_] glycerol (99%) (Cambridge Isotope Laboratories, Inc., Andover, MA, USA) and naturally labelled glycerol. Naturally labelled acetate was substituted with 99% [1-^13^C] sodium acetate, [2-^13^C] sodium acetate or a molar 1:1 mixture of [U-^13^C_2_] sodium acetate obtained from Sigma-Aldrich (Poole, Dorset, UK) and natural sodium acetate.

Starter cultures were prepared by inoculating LB medium with a loop of freshly plated PAO1. After 6 hr of incubation, 50 µL of cell suspension was transferred to a second culture of MOPS minimal medium. Subsequently, exponentially growing cells were used as inoculum for main cultures, and PAO1 was cultured in 25 mL of minimal medium in 250 mL baffled shake flasks (200 rpm, 37°C) in an orbital shaker (Aquatron, Infors AG, Switzerland). These growth conditions were selected to ensure sufficient aeration during cultivation. As shown previously for *P. aeruginosa* PAO1, the oxygen level was maintained above 80% of saturation under identical growth conditions [21]. In cultures incubated with ^13^C-tracer, the inoculum level was always kept below 1% (initial OD < 0.02) of the final sampled cell concentration to exclude potential interference of non-labelled inoculum on subsequent calculation of flux [94].

Mass isotopomer labelling analysis of proteinogenic amino acids, mass isotopomer labelling analysis of cell sugar monomers (glucose and glucosamine), metabolic reaction network and flux calculation were carried out as described in [25].

### Quantification of substrates and products

Acetate and glycerol, as well as organic acids (citric acid, α-ketoglutaric acid, gluconic acid, 2-ketogluconic acid, pyruvic acid, succinic acid, lactic acid, formic acid, fumaric acid) were quantified in filtered culture supernatants (Costar^®^ Spin-X^®^ 0.22 μm) using isocratic high-performance liquid chromatography (Agilent 1260 Infinity series, Aminex HPX-87H column at 65°C and a flow rate of 0.5 mL min^-1^) equipped with RI and UV detector (210 nm) with 50 mM H_2_SO_4_ as eluent [95]. Concentrations were determined from commercial standards which were analysed on the same run. These data were then used to calculate specific uptake and formation rates and yields for acetate, glycerol and secreted by-products, respectively (File S2).

### Calculation of redox cofactor and ATP balances

#### NADPH, NADH and FADH_2_

Total production of reduced cofactors was determined by summing up all cofactor-forming fluxes considering substrate-dependent cofactor specificities [30,96,97]. Anabolic NADPH requirements and anabolically produced NADH were estimated from the biomass composition [37,98] and measured specific growth rates. Surplus NADPH was considered to be converted into NADH *via* the activities of a soluble (SthA, PA2991) and a membrane-bound, proton-translocating (PntAB, PA0195-PA0196) pyridine nucleotide transhydrogenases [48].

#### ATP

Production of ATP *via* substrate-level phosphorylation was calculated by adding all ATP-producing fluxes and subtracting all ATP-consuming fluxes. Acetate uptake was considered *via* acetyl-CoA synthase (ACS) consuming 2 mol of ATP per mol acetate catabolized [58]. The ATP synthesized from NADH and FADH_2_ *via* oxidative phosphorylation in the respiratory chain was estimated assuming a P/O ratio of 1.875 for NADH [48,98,99] and 1.0 for FADH_2_ [100], respectively. The ATP demand was calculated by adding up the requirements for anabolism estimated from biomass composition, observed specific growth rates, and non-growth associated maintenance (NGAM) needs [48,98,99]. ATP surplus represents available ATP to fulfil growth-associated maintenance and other cellular ATP-consuming tasks.

#### NAD(P)(H) extraction

*P. aeruginosa* PAO1 cultures were grown in MOPS media containing a single carbon source (40 mM acetate, 15 mM glucose, 30 mM glycerol, or 30 mM succinate) at 37°C with shaking at 250 rpm, using a culture volume of 150 mL in a 2 L baffled Erlenmeyer flask. For each NAD(P)(H) extraction, 1.8 mL of culture was removed and immediately added to 7.5 mL ice-cold 100% methanol followed by centrifugation at 3220 × *g* for 14 min at 4°C to obtain a cell pellet. The pellet was resuspended in 0.2 M HCl for NAD(P)^+^ or 0.2 M NaOH for NAD(P)H extraction, before incubation at 52.5°C for 10 min followed by incubation on ice for 5 min. HCl or NaOH was then neutralised by the dropwise addition of 0.1 M NaOH or 0.1 M HCl, respectively, whilst vertexing at low speed. The mixture was then centrifuged for 5 min at 15,800 × *g* and 135 µL of the supernatant was removed for immediate NAD(P)(H) measurement or storage at −80°C. Samples were stored for a maximum of 1 week before measurement.

#### NAD(P)(H) measurement

NAD(P)(H) concentrations were measured using an enzyme cycling assay in a 96-well microtiter plate (Thermo Scientific 167008) as described in [101]. A reagent master mix was prepared containing 2 volumes 1 M bicine (pH 8.0), 1 volume 100% ethanol, 1 volume 40 mM EDTA (pH 8.0), 1 volume 4.2 mM thiazolyl blue, 2 volumes 16 mM phenazine ethosulfate, and 1 volume dH_2_O. The reagent mix was incubated at 30°C and primed to injectors in a BMG LABTECH FLUOstar Omega microplate reader. Aliquots (15 µL) of NAD(P)(H) extracts were added to individual wells of a 96-well microtiter plate, which was then incubated in the microplate reader at 30°C. Reagent master mix (80 µL) was added *via* the microplate reader injector (300 µL s^-1^) and vigorously mixed (200 rpm, 3 sec) before static incubation for 10 min. Immediately before measurement, a solution of alcohol dehydrogenase (1 mg mL^-1^ in 0.1 M bicine) was prepared for NAD(H) measurement or glucose 6-phosphate dehydrogenase (0.1 mg mL^-1^ in 0.1 M bicine) for NAD(P)(H) measurement and primed to a second injector. To start the reaction each well was injected (300 µL s^-1^) with 5 µL of enzyme solution, followed by vigorous mixing (200 rpm, 1 sec). The absorbance at 570 nm was then recorded every 30-60 sec for 20 min, with vigorous shaking (200 rpm, 1 sec) before each read. Slopes from plots of absorbance over time were calculated for NAD(P)H and NAD(P)^+^, which were then used to calculate ratios.

#### Colony Forming Unit (CFU) enumeration

Alongside each NAD(P)(H) extraction, an aliquot from the same culture was removed, serial diluted and plated onto LB agar using the single plate-serial dilution spotting method described in [102], and colonies were then grown overnight at 37°C.

#### Western-blot analysis

The cultures were grown aerobically to an OD_600_ of 0.5 in MOPS minimal medium supplemented with the indicated carbon sources. The samples were centrifuged at 3220 × *g* for 10 min. Equal amounts of protein were resolved on a 12% SDS—polyacrylamide gel. The proteins were blotted onto a nitrocellulose membrane, which was blocked with 5% (w/v) dried skimmed milk in TBS buffer. The membranes were probed with mouse anti-NirS [103] and with a IRDye^®^ 680RD and IRDye^®^ 800CW Goat anti-mouse IgG secondary antibodies (925-68070). Bands were visualized on an Odyssey Infrared Imaging System (LI-COR Biosciences.

#### Fluorescence microscopy

Fluorescence microscopy experiments to determine bacteria length were performed on a custom-built microscope based on an Olympus (Center Valley, PA) IX-73 frame with a 100 × 1.49 NA oil objective lens (Olympus UAPON100XOTIRF) and a 488-nm laser (Coherent Sapphire 488-300 CW CDRH). Samples were imaged using epi-illumination and images were relayed onto an Andor iXon Ultra 897 camera by a 1.3 × magnification Cairn Twincam image splitter (the second port of the image splitter was not used during these experiments). The resulting pixel width on the sample was measured to be 118 nm. For each field of interest, 100 frames at 100 msec exposure time were captured in a 256 × 256 pixel^2^ region.

#### Bacterial size analysis

The micrographs captured for this study exhibited typically one or two bacteria over a 30 mm x 30 mm^2^ area. The raw data were segmented and filtered for analysis using a MATLAB script included in the Supplementary Information. The data were first segmented: the MATLAB function adaptthresh was used to binarize the data using an adaptive threshold. This was deemed good enough as a segmentation step, since the bacteria were sparsely distributed over a dark background in a single frame. The MATLAB regionprops function was used to extract the area, major axis length (length of the major axis of a fitted ellipse to the detected blob shape), eccentricity, and centroid of each segmented blob. A filtering step was implemented to only include detected shapes with major axis lengths between 1.7 and 4.5 µm (14 and 38 pixels), corresponding to the possible size ranges for the bacteria.

## Acknowledgements

This work was funded by a grant (BB/M019411/1) awarded to MW from the BBSRC. ST was supported by a BBSRC DTP studentship. MK and CW acknowledge support by the German Federal Ministry for Education and Research (BMBF) through the grants “BioNylon” (FKZ 03V0757) and “LignoValue” (FKZ 031B0344A). SKD is currently supported by a Herchel Smith Postdoctoral Fellowship. Some elements of this work were supported by an EMBO Short Term Fellowship to SKD (7293-2017). CFK acknowledges funding from the UK Engineering and Physical Sciences Research Council, EPSRC (grants EP/L015889/1 and EP/H018301/1), the Wellcome Trust (grants 3-3249/Z/16/Z and 089703/Z/09/Z) and the UK Medical Research Council, MRC (grants MR/K015850/1 and MR/K02292X/1). PVF is supported by the Gates Cambridge Trust. We are grateful to Mrs. S. Buchmaier and Dr. G. Layer for providing the anti-NirS antibodies. pMF230 was a gift from Michael Franklin (Addgene plasmid # 62546; http://n2t.net/addgene:62546; RRID:Addgene_62546). We thank the Cambridge Centre for Proteomics, including Dr Mike Deery, Mrs. Renata Feret and Prof. Kathryn Lilley for proteomics support.

## Figure Legends

**Figure S1;** Transcriptomic analysis (CummeRbund) and proteomic analysis (LIMMA Package) of MOPS-acetate *versus* MOPS-glycerol grown *P. aeruginosa*. A; Principal component analysis (PCAplot) of RNA-seq data. BS; Scatterplot matrix of gene and isoform level expression (csScatterMatrix). C; A scatter plot comparing the FPKM values from the two conditions (csScatter). E; Heatmap and hierarchical clustering of proteomic data. F; Principal component analysis of proteomic data. G; Pairwise comparison of proteomic data.

**Figure S2;** NADH:NAD^+^ ratios alongside colony forming units (CFUs) from cells grown in MOPS A; Acetate, B; Glycerol, C: Glucose, D; Succinate. NADPH:NADP^+^ ratios alongside colony forming units (CFUs) from cells grown in MOPS E; Acetate, F; Glycerol, G: Glucose, H; Succinate. Total NADP(H) concentrations detected over multiple time points from cells grown in MOPS I; Acetate, J; Glycerol, K: Glucose, L: Succinate. Total NAD(H) concentrations detected over multiple time points from cells grown in MOPS M; Acetate, N; Glycerol, O: Glucose, P: Succinate. Three biological replicates per time-point.

**Figure S3;** Luciferase detection in *P. aeruginosa* PAO1 wild-type carrying chromosomal promoter: *lux* fusions for the above promoter regions, grown on various carbon sources, MOPS-acetate, MOPS-glucose, MOPS-glycerol, MOPS-succinate. A; *aceA*, B; *glcB*, C; *cco1*, D; *cco2*, E; *cox*, F; *cyo*, G; *glpD*, H; *nir*. Values normalised to OD_600_ (RLU/OD_600_). Three biological replicates per sample. Data analysed in GraphPad Prism (V 6.01) using t-test statistical analysis (MOPS glycerol *versus* acetate, glucose or succinate).

**Figure S4;** A; Colony forming units (CFUs) B; NAD(H) concentrations, and C: NADP(H) concentrations extracted from PAO1 wild-type grown in MOPS-acetate -/+ 20 mM KNO_3_. Three biological replicates per time-point. D; Colony forming units (CFUs) E; NAD(H) concentrations, and F: NADP(H) concentrations extracted from PAO1 WT and Δ*dnr* cells grown cells grown in MOPS-acetate -/+ 20 mM KNO_3_. Three biological replicates per time-point.

**Figure S5;** Western blot showing; (A) Expression of the Anr/Dnr-regulated denitrification enzyme NirS during exponential growth of *P. aeruginosa* in MOPS-glucose, MOPS-succinate and MOPS-acetate. NirS is not expressed in a Δ*anr* or Δ*dnr* mutant. The isocitrate dehydrogenases (ICD and IDH) were used as loading controls. (B) Expression of NirS during exponential growth of *P. aeruginosa* in MOPS-acetate and MOPS-acetate + sodium nitrate (20 mM). Sodium nitrate (20 mM) is capable of inducing NirS expression in the WT and in a Δr*oxSR* mutant, but not in a Δ*anr* or Δ*dnr* mutant. Three biological replicates per sample. The Δ*anr*, Δ*dnr* and Δ*roxR* lanes are representative of the triplicates.

**Figure S6;** Bacterial cell length as determined by fluorescence microscopy, PAO1 carrying the pMF230 eGFP-expressing plasmid. Cells grown in MOPS media with acetate, glucose, glycerol or succinate as the sole carbon sources. Actual data points shown in File S3.

**Table S1;** A; Oligonucleotide primers used in this study. B; Bacterial strains and plasmids used in this study [14,51,84,104].

**File_S1;** Full transcriptomics (tab A) and proteomics (tab B) of MOPS-acetate *versus* MOPS-glycerol grown *P. aeruginosa*.

**File_S2;** ^13^C fluxomics data of MOPS acetate and MOPS glycerol grown *P. aeruginosa*, including calculations for anabolic demand (A),OpenFLUX SimVector files (B), reaction network (C), goodness of fit (D) and metabolic fluxes (E).

**File_S3;** Correlation plot calculations to illustrate the log2 fold-change differences between the transcriptomic, proteomic and fluxomic data (A-C) and cell size data (D).

## References

1. Klockgether J, Tümmler B. Recent advances in understanding Pseudomonas aeruginosa as a pathogen. F1000Research. 2017;6: 1261. doi:10.12688/f1000research.10506.1

2. Rajan S, Saiman L. Pulmonary infections in patients with cystic fibrosis. Semin Respir Infect. 2002;17: 47–56. Available: http://www.ncbi.nlm.nih.gov/pubmed/11891518

3. Gibson RL, Burns JL, Ramsey BW. Pathophysiology and Management of Pulmonary Infections in Cystic Fibrosis. Am J Respir Crit Care Med. 2003;168: 918–951. doi:10.1164/rccm.200304-505SO

4. La Rosa R, Johansen HK, Molin S. Convergent Metabolic Specialization through Distinct Evolutionary Paths in Pseudomonas aeruginosa. MBio. American Society for Microbiology; 2018;9: e00269–18. doi:10.1128/mBio.00269-18

5. Rojo F. Carbon catabolite repression in Pseudomonas⍰: optimizing metabolic versatility and interactions with the environment. FEMS Microbiol Rev. 2010;34: 658–684. doi:10.1111/j.1574-6976.2010.00218.x

6. Palmer KL, Aye LM, Whiteley M. Nutritional Cues Control Pseudomonas aeruginosa Multicellular Behavior in Cystic Fibrosis Sputum. J Bacteriol. 2007;189: 8079–8087. doi:10.1128/JB.01138-07

7. Flynn JM, Niccum D, Dunitz JM, Hunter RC. Evidence and Role for Bacterial Mucin Degradation in Cystic Fibrosis Airway Disease. Wozniak DJ, editor. PLOS Pathog. Public Library of Science; 2016;12: e1005846. doi:10.1371/journal.ppat.1005846

8. Sun Z, Kang Y, Norris MH, Troyer RM, Son MS, Schweizer HP, et al. Blocking Phosphatidylcholine Utilization in Pseudomonas aeruginosa, via Mutagenesis of Fatty Acid, Glycerol and Choline Degradation Pathways, Confirms the Importance of This Nutrient Source In Vivo. van Veen HW, editor. PLoS One. Public Library of Science; 2014;9: e103778. doi:10.1371/journal.pone.0103778

9. Marty N, Dournes JL, Chabanon G, Montrozier H. Influence of nutrient media on the chemical composition of the exopolysaccharide from mucoid and non-mucoid Pseudomonas aeruginosa. FEMS Microbiol Lett. 1992;77: 35–44. Available: http://www.ncbi.nlm.nih.gov/pubmed/1459419

10. Kim J, Oliveros JC, Nikel PI, de Lorenzo V, Silva-Rocha R. Transcriptomic fingerprinting of Pseudomonas putida under alternative physiological regimes. Environ Microbiol Rep. 2013;5: 883–891. doi:10.1111/1758-2229.12090

11. Nikel PI, Romero-Campero FJ, Zeidman JA, Goñi-Moreno Á, de Lorenzo V. The glycerol-dependent metabolic persistence of Pseudomonas putida KT2440 reflects the regulatory logic of the GlpR repressor. MBio. American Society for Microbiology; 2015;6: e00340–15. doi:10.1128/mBio.00340-15

12. Shuman J, Giles TX, Carroll L, Tabata K, Powers A, Suh S-J, et al. Transcriptome analysis of a Pseudomonas aeruginosa sn-glycerol-3-phosphate dehydrogenase mutant reveals a disruption in bioenergetics. Microbiology. Microbiology Society; 2018;164: 551–562. doi:10.1099/mic.0.000646

13. Agrawal S, Jaswal K, Shiver AL, Balecha H, Patra T, Chaba R. A genome-wide screen in Escherichia coli reveals that ubiquinone is a key antioxidant for metabolism of long-chain fatty acids. J Biol Chem. 2017;292: 20086–20099. doi:10.1074/jbc.M117.806240

14. Choi K-H, Schweizer HP. mini-Tn7 insertion in bacteria with single attTn7 sites: example Pseudomonas aeruginosa. Nat Protoc. Nature Publishing Group; 2006;1: 153–161. doi:10.1038/nprot.2006.24

15. Liebermeister W, Noor E, Flamholz A, Davidi D, Bernhardt J, Milo R. Visual account of protein investment in cellular functions. Proc Natl Acad Sci U S A. National Academy of Sciences; 2014;111: 8488–93. doi:10.1073/pnas.1314810111

16. Beste DJ V, Nöh K, Niedenführ S, Mendum TA, Hawkins ND, Ward JL, et al. 13C-flux spectral analysis of host-pathogen metabolism reveals a mixed diet for intracellular Mycobacterium tuberculosis. Chem Biol. Elsevier; 2013;20: 1012–21. doi:10.1016/j.chembiol.2013.06.012

17. Eisenreich W, Slaghuis J, Laupitz R, Bussemer J, Stritzker J, Schwarz C, et al. 13 C isotopologue perturbation studies of Listeria monocytogenes carbon metabolism and its modulation by the virulence regulator PrfA. Proc Natl Acad Sci. 2006;103: 2040–2045. doi:10.1073/pnas.0507580103

18. Long CP, Gonzalez JE, Feist AM, Palsson BO, Antoniewicz MR. Dissecting the genetic and metabolic mechanisms of adaptation to the knockout of a major metabolic enzyme in Escherichia coli. Proc Natl Acad Sci U S A. National Academy of Sciences; 2018;115: 222–227. doi:10.1073/pnas.1716056115

19. Lien SK, Niedenführ S, Sletta H, Nöh K, Bruheim P. Fluxome study of Pseudomonas fluorescens reveals major reorganisation of carbon flux through central metabolic pathways in response to inactivation of the anti-sigma factor MucA. BMC Syst Biol. BioMed Central; 2015;9: 6. doi:10.1186/s12918-015-0148-0

20. Lassek C, Berger A, Zühlke D, Wittmann C, Riedel K. Proteome and carbon flux analysis of Pseudomonas aeruginosa clinical isolates from different infection sites. Proteomics. Wiley-Blackwell; 2016;16: 1381–1385. doi:10.1002/pmic.201500228

21. Berger A, Dohnt K, Tielen P, Jahn D, Becker J, Wittmann C. Robustness and plasticity of metabolic pathway flux among uropathogenic isolates of Pseudomonas aeruginosa. Fong SS, editor. PLoS One. Public Library of Science; 2014;9: e88368. doi:10.1371/journal.pone.0088368

22. Opperman MJ, Shachar-Hill Y. Metabolic flux analyses of Pseudomonas aeruginosa cystic fibrosis isolates. Metab Eng. Academic Press; 2016;38: 251–263. doi:10.1016/J.YMBEN.2016.09.002

23. Collier DN, Hager PW, Phibbs P V. Catabolite repression control in the Pseudomonads. Res Microbiol. 147: 551–61. Available: http://www.ncbi.nlm.nih.gov/pubmed/9084769

24. Rossi E, Falcone M, Molin S, Johansen HK. High-resolution in situ transcriptomics of Pseudomonas aeruginosa unveils genotype independent patho-phenotypes in cystic fibrosis lungs. Nat Commun. Nature Publishing Group; 2018;9: 3459. doi:10.1038/s41467-018-05944-5

25. Kohlstedt M, Wittmann C. GC-MS-based 13C metabolic flux analysis resolves the parallel and cyclic glucose metabolism of Pseudomonas putida KT2440 and Pseudomonas aeruginosa PAO1. Metab Eng. 2019;54: 35–53. doi:10.1016/j.ymben.2019.01.008

26. Schweizer HP, Po C. Regulation of glycerol metabolism in Pseudomonas aeruginosa: characterization of the glpR repressor gene. J Bacteriol. American Society for Microbiology Journals; 1996;178: 5215–21. doi:10.1128/JB.178.17.5215-5221.1996

27. Williams SG, Greenwood JA, Jones CW. The effect of nutrient limitation on glycerol uptake and metabolism in continuous cultures of Pseudornonas aeruginosa [Internet]. 2019. Available: http://www.microbiologyresearch.org

28. Poblete-Castro I, Wittmann C, Nikel PI. Biochemistry, genetics and biotechnology of glycerol utilization in Pseudomonas species. Microb Biotechnol. 2019; 1751-7915.13400. doi:10.1111/1751-7915.13400

29. Conway T. The Entner-Doudoroff pathway: history, physiology and molecular biology. FEMS Microbiol Lett. No longer published by Elsevier; 1992;103: 1–27. doi:10.1016/0378-1097(92)90334-K

30. Nikel PI, Chavarría M, Fuhrer T, Sauer U, de Lorenzo V. Pseudomonas putida KT2440 Strain Metabolizes Glucose through a Cycle Formed by Enzymes of the Entner-Doudoroff, Embden-Meyerhof-Parnas, and Pentose Phosphate Pathways. J Biol Chem. American Society for Biochemistry and Molecular Biology; 2015;290: 25920–32. doi:10.1074/jbc.M115.687749

31. Heath, H. E., Elizabeth GT. Relationship Between Catabolism of Glycerol and Metabolism of Hexosephosphate Derivatives by Pseudomonas aeruginosa [Internet]. JOURNAL OF BACTERIOLOGY. 1978. Available: http://jb.asm.org/

32. Diaz-Perez AL, Roman-Doval C, Diaz-Perez C, Cervantes C, Sosa-Aguirre CR, Lopez-Meza JE, et al. Identification of the aceA gene encoding isocitrate lyase required for the growth of Pseudomonas aeruginosa on acetate, acyclic terpenes and leucine. FEMS Microbiol Lett. 2007;269: 309–316. doi:10.1111/j.1574-6968.2007.00654.x

33. McVey AC, Medarametla P, Chee X, Bartlett S, Poso A, Spring DR, et al. Structural and Functional Characterization of Malate Synthase G from Opportunistic Pathogen Pseudomonas aeruginosa. Biochemistry. American Chemical Society; 2017;56: 5539–5549. doi:10.1021/acs.biochem.7b00852

34. Jacob K, Rasmussen A, Tyler P, Servos MM, Sylla M, Prado C, et al. Regulation of acetyl-CoA synthetase transcription by the CrbS/R two-component system is conserved in genetically diverse environmental pathogens. PLoS One. Public Library of Science; 2017;12: e0177825. doi:10.1371/journal.pone.0177825

35. Görisch H, Jeoung J-H, Rückert A, Kretzschmar U. Malate:quinone oxidoreductase is essential for growth on ethanol or acetate in Pseudomonas aeruginosa. Microbiology. 2002;148: 3839–3847. doi:10.1099/00221287-148-12-3839

36. Crousilles A, Dolan SK, Brear P, Chirgadze DY, Welch M. Gluconeogenic precursor availability regulates flux through the glyoxylate shunt in Pseudomonas aeruginosa. J Biol Chem. American Society for Biochemistry and Molecular Biology; 2018;293: 14260–14269. doi:10.1074/jbc.RA118.004514

37. Berger A, Dohnt K, Tielen P, Jahn D, Becker J, Wittmann C. Robustness and Plasticity of Metabolic Pathway Flux among Uropathogenic Isolates of Pseudomonas aeruginosa. Fong SS, editor. PLoS One. Public Library of Science; 2014;9: e88368. doi:10.1371/journal.pone.0088368

38. Singh R, Mailloux RJ, Puiseux-Dao S, Appanna VD. Oxidative stress evokes a metabolic adaptation that favors increased NADPH synthesis and decreased NADH production in Pseudomonas fluorescens. J Bacteriol. 2007;189: 6665–6675. doi:10.1128/JB.00555-07

39. Arai H, Kawakami T, Osamura T, Hirai T, Sakai Y, Ishii M. Enzymatic characterization and in vivo function of five terminal oxidases in Pseudomonas aeruginosa. J Bacteriol. American Society for Microbiology Journals; 2014;196: 4206–15. doi:10.1128/JB.02176-14

40. Dietrich LEP, Okegbe C, Price-Whelan A, Sakhtah H, Hunter RC, Newman DK. Bacterial community morphogenesis is intimately linked to the intracellular redox state. J Bacteriol. American Society for Microbiology Journals; 2013;195: 1371–80. doi:10.1128/JB.02273-12

41. Poole RK, Cook GM. Redundancy of aerobic respiratory chains in bacteria? Routes, reasons and regulation. Adv Microb Physiol. 2000;43: 165–224. Available: http://www.ncbi.nlm.nih.gov/pubmed/10907557

42. Raba DA, Rosas-Lemus M, Menzer WM, Li C, Fang X, Liang P, et al. Characterization of the Pseudomonas aeruginosa NQR Complex, a Bacterial Proton Pump with Roles in Autopoisoning Resistance. J Biol Chem. American Society for Biochemistry and Molecular Biology; 2018; jbc.RA118.003194. doi:10.1074/jbc.RA118.003194

43. Arai H. Regulation and Function of Versatile Aerobic and Anaerobic Respiratory Metabolism in Pseudomonas aeruginosa. Front Microbiol. Frontiers Media SA; 2011;2: 103. doi:10.3389/fmicb.2011.00103

44. Comolli JC, Donohue TJ. Differences in two Pseudomonas aeruginosa cbb3 cytochrome oxidases. Mol Microbiol. Wiley/Blackwell (10.1111); 2004;51: 1193–1203. doi:10.1046/j.1365-2958.2003.03904.x

45. Le Laz S, kpebe A, Bauzan M, Lignon S, Rousset M, Brugna M. Expression of terminal oxidases under nutrient-starved conditions in Shewanella oneidensis: detection of the A-type cytochrome c oxidase. Sci Rep. Nature Publishing Group; 2016;6: 19726. doi:10.1038/srep19726

46. Osamura T, Kawakami T, Kido R, Ishii M, Arai H. Specific expression and function of the A-type cytochrome c oxidase under starvation conditions in Pseudomonas aeruginosa. PLoS One. Public Library of Science; 2017;12: e0177957. doi:10.1371/journal.pone.0177957

47. Kawakami T, Kuroki M, Ishii M, Igarashi Y, Arai H. Differential expression of multiple terminal oxidases for aerobic respiration in Pseudomonas aeruginosa. Environ Microbiol. Wiley/Blackwell (10.1111); 2009;12: 1399–1412. doi:10.1111/j.1462-2920.2009.02109.x

48. Nikel PI, Pérez-Pantoja D, de Lorenzo V. Pyridine nucleotide transhydrogenases enable redox balance of Pseudomonas putida during biodegradation of aromatic compounds. Environ Microbiol. Wiley/Blackwell (10.1111); 2016;18: 3565–3582. doi:10.1111/1462-2920.13434

49. Sauer U, Canonaco F, Heri S, Perrenoud A, Fischer E. The Soluble and Membrane-bound Transhydrogenases UdhA and PntAB Have Divergent Functions in NADPH Metabolism of Escherichia coli. J Biol Chem. 2004;279: 6613–6619. doi:10.1074/jbc.M311657200

50. Szenk M, Dill KA, de Graff AMR. Why Do Fast-Growing Bacteria Enter Overflow Metabolism? Testing the Membrane Real Estate Hypothesis. Cell Syst. 2017;5: 95–104. doi:10.1016/j.cels.2017.06.005

51. Nivens DE, Ohman DE, Williams J, Franklin MJ. Role of Alginate and Its O Acetylation in Formation of Pseudomonas aeruginosa Microcolonies and Biofilms. J Bacteriol. 2001;183: 1047–1057. doi:10.1128/JB.183.3.1047-1057.2001

52. Deforet M, van Ditmarsch D, Xavier JB. Cell-Size Homeostasis and the Incremental Rule in a Bacterial Pathogen. Biophys J. The Biophysical Society; 2015;109: 521–8. doi:10.1016/j.bpj.2015.07.002

53. Borrero-de Acuña JM, Rohde M, Wissing J, Jänsch L, Schobert M, Molinari G, et al. Protein Network of the Pseudomonas aeruginosa Denitrification Apparatus. J Bacteriol. American Society for Microbiology Journals; 2016;198: 1401–13. doi:10.1128/JB.00055-16

54. Line L, Alhede M, Kolpen M, KÃ¼hl M, Ciofu O, Bjarnsholt T, et al. Physiological levels of nitrate support anoxic growth by denitrification of Pseudomonas aeruginosa at growth rates reported in cystic fibrosis lungs and sputum. Front Microbiol. Frontiers; 2014;5: 554. doi:10.3389/fmicb.2014.00554

55. Worlitzsch D, Tarran R, Ulrich M, Schwab U, Cekici A, Meyer KC, et al. Effects of reduced mucus oxygen concentration in airway Pseudomonas infections of cystic fibrosis patients. J Clin Invest. American Society for Clinical Investigation; 2002;109: 317. doi:10.1172/JCI13870

56. Palmer KL, Brown SA, Whiteley M. Membrane-bound nitrate reductase is required for anaerobic growth in cystic fibrosis sputum. J Bacteriol. American Society for Microbiology Journals; 2007;189: 4449–55. doi:10.1128/JB.00162-07

57. Chen J, Strous M. Denitrification and aerobic respiration, hybrid electron transport chains and co-evolution. Biochim Biophys Acta - Bioenerg. Elsevier; 2013;1827: 136–144. doi:10.1016/J.BBABIO.2012.10.002

58. Görisch H, Kretzschmar U, Schobert M. The Pseudomonas aeruginosa acsA gene, encoding an acetyl-CoA synthetase, is essential for growth on ethanol. Microbiology. 2001;147: 2671–2677. doi:10.1099/00221287-147-10-2671

59. Vemuri GN, Eiteman MA, McEwen JE, Olsson L, Nielsen J. Increasing NADH oxidation reduces overflow metabolism in Saccharomyces cerevisiae. Proc Natl Acad Sci U S A. National Academy of Sciences; 2007;104: 2402–7. doi:10.1073/pnas.0607469104

60. Dietrich LEP, Okegbe C, Price-Whelan A, Sakhtah H, Hunter RC, Newman DK. Bacterial community morphogenesis is intimately linked to the intracellular redox state. J Bacteriol. American Society for Microbiology Journals; 2013;195: 1371–80. doi:10.1128/JB.02273-12

61. Ishii N, Nakahigashi K, Baba T, Robert M, Soga T, Kanai A, et al. Multiple High-Throughput Analyses Monitor the Response of E. coli to Perturbations. Science (80-). 2007;316: 593–597. doi:10.1126/science.1132067

62. Buschke N, Becker J, Schäfer R, Kiefer P, Biedendieck R, Wittmann C. Systems metabolic engineering of xylose-utilizing Corynebacterium glutamicum for production of 1,5-diaminopentane. Biotechnol J. 2013;8: 557–570. doi:10.1002/biot.201200367

63. Kohlstedt M, Sappa PK, Meyer H, Maaß S, Zaprasis A, Hoffmann T, et al. Adaptation of Bacillus subtilis carbon core metabolism to simultaneous nutrient limitation and osmotic challenge: a multi-omics perspective. Environ Microbiol. 2014;16: 1898–1917. doi:10.1111/1462-2920.12438

64. Moxley JF, Jewett MC, Antoniewicz MR, Villas-Boas SG, Alper H, Wheeler RT, et al. Linking high-resolution metabolic flux phenotypes and transcriptional regulation in yeast modulated by the global regulator Gcn4p. Proc Natl Acad Sci. 2009;106: 6477–6482. doi:10.1073/pnas.0811091106

65. Lange A, Becker J, Schulze D, Cahoreau E, Portais J-C, Haefner S, et al. Bio-based succinate from sucrose: High-resolution 13C metabolic flux analysis and metabolic engineering of the rumen bacterium Basfia succiniciproducens. Metab Eng. 2017;44: 198–212. doi:10.1016/j.ymben.2017.10.003

66. Kukurugya MA, Mendonca CM, Solhtalab M, Wilkes RA, Thannhauser TW, Aristilde L. Multi-omics analysis unravels a segregated metabolic flux network that tunes co-utilization of sugar and aromatic carbons in Pseudomonas putida. J Biol Chem. American Society for Biochemistry and Molecular Biology; 2019;294: 8464–8479. doi:10.1074/jbc.RA119.007885

67. Fuhrer T, Fischer E, Sauer U. Experimental identification and quantification of glucose metabolism in seven bacterial species. J Bacteriol. American Society for Microbiology Journals; 2005;187: 1581–90. doi:10.1128/JB.187.5.1581-1590.2005

68. Kwon T, Huse HK, Vogel C, Whiteley M, Marcotte EM. Protein-to-mRNA ratios are conserved between Pseudomonas aeruginosa strains. J Proteome Res. American Chemical Society; 2014;13: 2370–80. doi:10.1021/pr4011684

69. Nikel PI, Kim J, de Lorenzo V. Metabolic and regulatory rearrangements underlying glycerol metabolism in Pseudomonas putida KT2440. Environ Microbiol. John Wiley & Sons, Ltd (10.1111); 2014;16: 239–254. doi:10.1111/1462-2920.12224

70. Cattoir V, Narasimhan G, Skurnik D, Aschard H, Roux D, Ramphal R, et al. Transcriptional response of mucoid Pseudomonas aeruginosa to human respiratory mucus. MBio. American Society for Microbiology (ASM); 2013;3: e00410–12. doi:10.1128/mBio.00410-12

71. Wu J, Bauer CE. RegB kinase activity is controlled in part by monitoring the ratio of oxidized to reduced ubiquinones in the ubiquinone pool. MBio. American Society for Microbiology; 2010;1: e00272–10. doi:10.1128/mBio.00272-10

72. Price-Whelan A, Dietrich LEP, Newman DK. Pyocyanin alters redox homeostasis and carbon flux through central metabolic pathways in Pseudomonas aeruginosa PA14. J Bacteriol. American Society for Microbiology Journals; 2007;189: 6372–81. doi:10.1128/JB.00505-07

73. Lin Y-C, Sekedat MD, Cornell WC, Silva GM, Okegbe C, Price-Whelan A, et al. Phenazines Regulate Nap-Dependent Denitrification in Pseudomonas aeruginosa Biofilms. J Bacteriol. American Society for Microbiology Journals; 2018;200: e00031–18. doi:10.1128/JB.00031-18

74. Lycus P, Soriano-Laguna MJ, Kjos M, Richardson DJ, Gates AJ, Milligan DA, et al. A bet-hedging strategy for denitrifying bacteria curtails their release of N2O. Proc Natl Acad Sci U S A. National Academy of Sciences; 2018;115: 11820–11825. doi:10.1073/pnas.1805000115

75. Alvarez-Ortega C, Harwood CS. Responses of Pseudomonas aeruginosa to low oxygen indicate that growth in the cystic fibrosis lung is by aerobic respiration. Mol Microbiol. Wiley/Blackwell (10.1111); 2007;65: 153–165. doi:10.1111/j.1365-2958.2007.05772.x

76. Chen J, Strous M. Denitrification and aerobic respiration, hybrid electron transport chains and co-evolution. Biochim Biophys Acta - Bioenerg. Elsevier; 2013;1827: 136–144. doi:10.1016/J.BBABIO.2012.10.002

77. Son MS, Matthews WJ, Kang Y, Nguyen DT, Hoang TT. In vivo evidence of Pseudomonas aeruginosa nutrient acquisition and pathogenesis in the lungs of cystic fibrosis patients. Infect Immun. 2007;75: 5313–5324. doi:10.1128/IAI.01807-06

78. Rossi E, Falcone M, Molin S, Johansen HK. High-resolution in situ transcriptomics of Pseudomonas aeruginosa unveils genotype independent patho-phenotypes in cystic fibrosis lungs. Nat Commun. Nature Publishing Group; 2018;9: 3459. doi:10.1038/s41467-018-05944-5

79. Cornforth DM, Dees JL, Ibberson CB, Huse HK, Mathiesen IH, Kirketerp-Møller K, et al. Pseudomonas aeruginosa transcriptome during human infection. Proc Natl Acad Sci. 2018;115: E5125–E5134. doi:10.1073/pnas.1717525115

80. Jackson AA, Daniels EF, Hammond JH, Willger SD, Hogan DA. Global regulator Anr represses PlcH phospholipase activity in Pseudomonas aeruginosa when oxygen is limiting. Microbiology. Microbiology Society; 2014;160: 2215–25. doi:10.1099/mic.0.081158-0

81. Dötsch A, Eckweiler D, Schniederjans M, Zimmermann A, Jensen V, Scharfe M, et al. The Pseudomonas aeruginosa Transcriptome in Planktonic Cultures and Static Biofilms Using RNA Sequencing. Semsey S, editor. PLoS One. Public Library of Science; 2012;7: e31092. doi:10.1371/journal.pone.0031092

82. Vemuri GN, Altman E, Sangurdekar DP, Khodursky AB, Eiteman MA. Overflow metabolism in Escherichia coli during steady-state growth: transcriptional regulation and effect of the redox ratio. Appl Environ Microbiol. American Society for Microbiology (ASM); 2006;72: 3653–61. doi:10.1128/AEM.72.5.3653-3661.2006

83. Basan M, Hui S, Okano H, Zhang Z, Shen Y, Williamson JR, et al. Overflow metabolism in Escherichia coli results from efficient proteome allocation. Nature. Nature Publishing Group; 2015;528: 99–104. doi:10.1038/nature15765

84. Stover CK, Pham XQ, Erwin AL, Mizoguchi SD, Warrener P, Hickey MJ, et al. Complete genome sequence of Pseudomonas aeruginosa PAO1, an opportunistic pathogen. Nature. Nature Publishing Group; 2000;406: 959–964. doi:10.1038/35023079

85. LaBauve AE, Wargo MJ. Growth and laboratory maintenance of Pseudomonas aeruginosa. Curr Protoc Microbiol. NIH Public Access; 2012;Chapter 6: Unit 6E.1. doi:10.1002/9780471729259.mc06e01s25

86. Tata M, Wolfinger MT, Amman F, Roschanski N, Dötsch A, Sonnleitner E, et al. RNASeq Based Transcriptional Profiling of Pseudomonas aeruginosa PA14 after Short- and Long-Term Anoxic Cultivation in Synthetic Cystic Fibrosis Sputum Medium. Roop RM, editor. PLoS One. Public Library of Science; 2016;11: e0147811. doi:10.1371/journal.pone.0147811

87. Bolger AM, Lohse M, Usadel B. Trimmomatic: a flexible trimmer for Illumina sequence data. Bioinformatics. 2014;30: 2114–2120. doi:10.1093/bioinformatics/btu170

88. Trapnell C, Roberts A, Goff L, Pertea G, Kim D, Kelley DR, et al. Differential gene and transcript expression analysis of RNA-seq experiments with TopHat and Cufflinks. Nat Protoc. NIH Public Access; 2012;7: 562–78. doi:10.1038/nprot.2012.016

89. McAlister GC, Nusinow DP, Jedrychowski MP, Wühr M, Huttlin EL, Erickson BK, et al. MultiNotch MS3 Enables Accurate, Sensitive, and Multiplexed Detection of Differential Expression across Cancer Cell Line Proteomes. Anal Chem. American Chemical Society; 2014;86: 7150–7158. doi:10.1021/ac502040v

90. Gatto L, Lilley KS. MSnbase-an R/Bioconductor package for isobaric tagged mass spectrometry data visualization, processing and quantitation. Bioinformatics. 2012;28: 288–289. doi:10.1093/bioinformatics/btr645

91. Smyth GK. limma: Linear Models for Microarray Data. Bioinformatics and Computational Biology Solutions Using R and Bioconductor. New York: Springer-Verlag; 2005. pp. 397–420. doi:10.1007/0-387-29362-0_23

92. Benjamini Y, Hochberg Y. Controlling the False Discovery Rate: A Practical and Powerful Approach to Multiple Testing [Internet]. Journal of the Royal Statistical Society. Series B (Methodological). WileyRoyal Statistical Society; 1995. pp. 289–300. doi:10.2307/2346101

93. Heath Damron F, McKenney ES, Barbier M, Liechti GW, Goldberg JB. Construction of Mobilizable Mini-Tn7 Vectors for Bioluminescent Detection and Single Copy Promoter lux Reporter Analysis in Gram-Negative Bacteria. CAMBRIDGE Univ Libr. 2013; doi:10.1128/AEM.00640-13

94. Wittmann C. Fluxome analysis using GC-MS. Microb Cell Fact. BioMed Central; 2007;6: 6. doi:10.1186/1475-2859-6-6

95. Kind S, Becker J, Wittmann C. Increased lysine production by flux coupling of the tricarboxylic acid cycle and the lysine biosynthetic pathway—Metabolic engineering of the availability of succinyl-CoA in Corynebacterium glutamicum. Metab Eng. 2013;15: 184–195. doi:10.1016/j.ymben.2012.07.005

96. Görisch H, Jeoung J-H, Rückert A, Kretzschmar U. Malate:quinone oxidoreductase is essential for growth on ethanol or acetate in Pseudomonas aeruginosa. Microbiology. 2002;148: 3839–3847. doi:10.1099/00221287-148-12-3839

97. Rivers DB, Blevins WT. Multiple Enzyme Forms of Glyceraldehyde-3-phosphate Dehydrogenase in Pseudomonas aeruginosa PAO. Microbiology. 1987;133: 3159–3164. doi:10.1099/00221287-133-11-3159

98. Bartell JA, Blazier AS, Yen P, Thøgersen JC, Jelsbak L, Goldberg JB, et al. Reconstruction of the metabolic network of Pseudomonas aeruginosa to interrogate virulence factor synthesis. Nat Commun. Nature Publishing Group; 2017;8: 14631. doi:10.1038/ncomms14631

99. Oberhardt MA, Puchałka J, Martins dos Santos VAP, Papin JA. Reconciliation of Genome-Scale Metabolic Reconstructions for Comparative Systems Analysis. Bourne PE, editor. PLoS Comput Biol. 2011;7: e1001116. doi:10.1371/journal.pcbi.1001116

100. Yuan Q, Huang T, Li P, Hao T, Li F, Ma H, et al. Pathway-Consensus Approach to Metabolic Network Reconstruction for Pseudomonas putida KT2440 by Systematic Comparison of Published Models. Virolle M-J, editor. PLoS One. Public Library of Science; 2017;12: e0169437. doi:10.1371/journal.pone.0169437

101. Kern SE, Price-Whelan A, Newman DK. Extraction and Measurement of NAD(P)+ and NAD(P)H. 2014. pp. 311–323. doi:10.1007/978-1-4939-0473-0_26

102. Thomas P, Sekhar AC, Upreti R, Mujawar MM, Pasha SS. Optimization of single plate-serial dilution spotting (SP-SDS) with sample anchoring as an assured method for bacterial and yeast cfu enumeration and single colony isolation from diverse samples. Biotechnol Reports. 2015;8: 45–55. doi:10.1016/j.btre.2015.08.003

103. Nicke T, Schnitzer T, Münch K, Adamczack J, Haufschildt K, Buchmeier S, et al. Maturation of the cytochrome cd1 nitrite reductase NirS from Pseudomonas aeruginosa requires transient interactions between the three proteins NirS, NirN and NirF. Biosci Rep. Portland Press Ltd; 2013;33. doi:10.1042/BSR20130043

104. Hoang TT, Karkhoff-Schweizer RR, Kutchma AJ, Schweizer HP. A broad-host-range Flp-FRT recombination system for site-specific excision of chromosomally-located DNA sequences: application for isolation of unmarked Pseudomonas aeruginosa mutants. Gene. 1998;212: 77–86. Available: http://www.ncbi.nlm.nih.gov/pubmed/9661666

