## Supplementary figures and images for "Contextual flexibility in *Pseudomonas aeruginosa* central carbon metabolism during growth in single carbon sources"

### Figure_S1

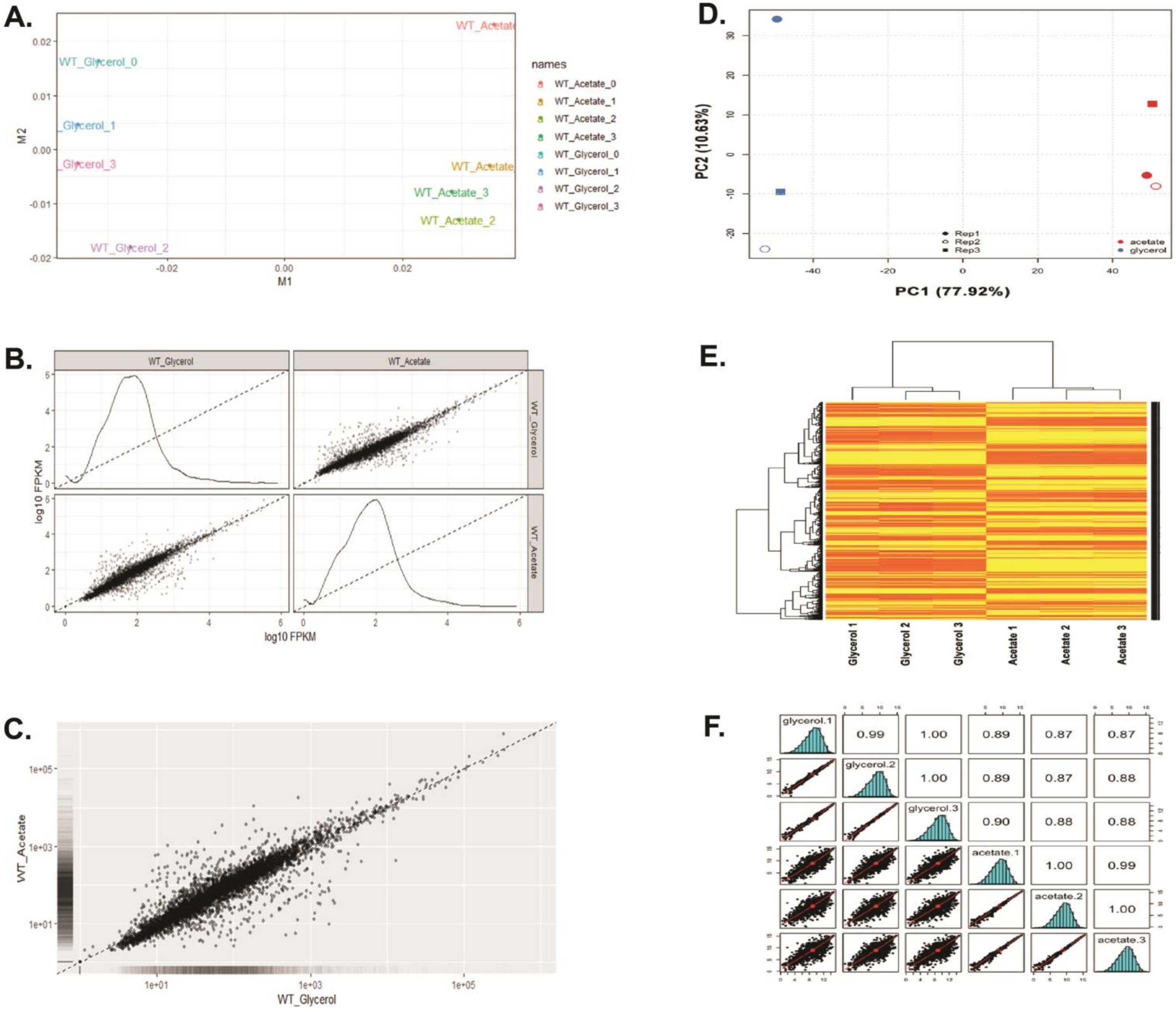

### Figure_S2

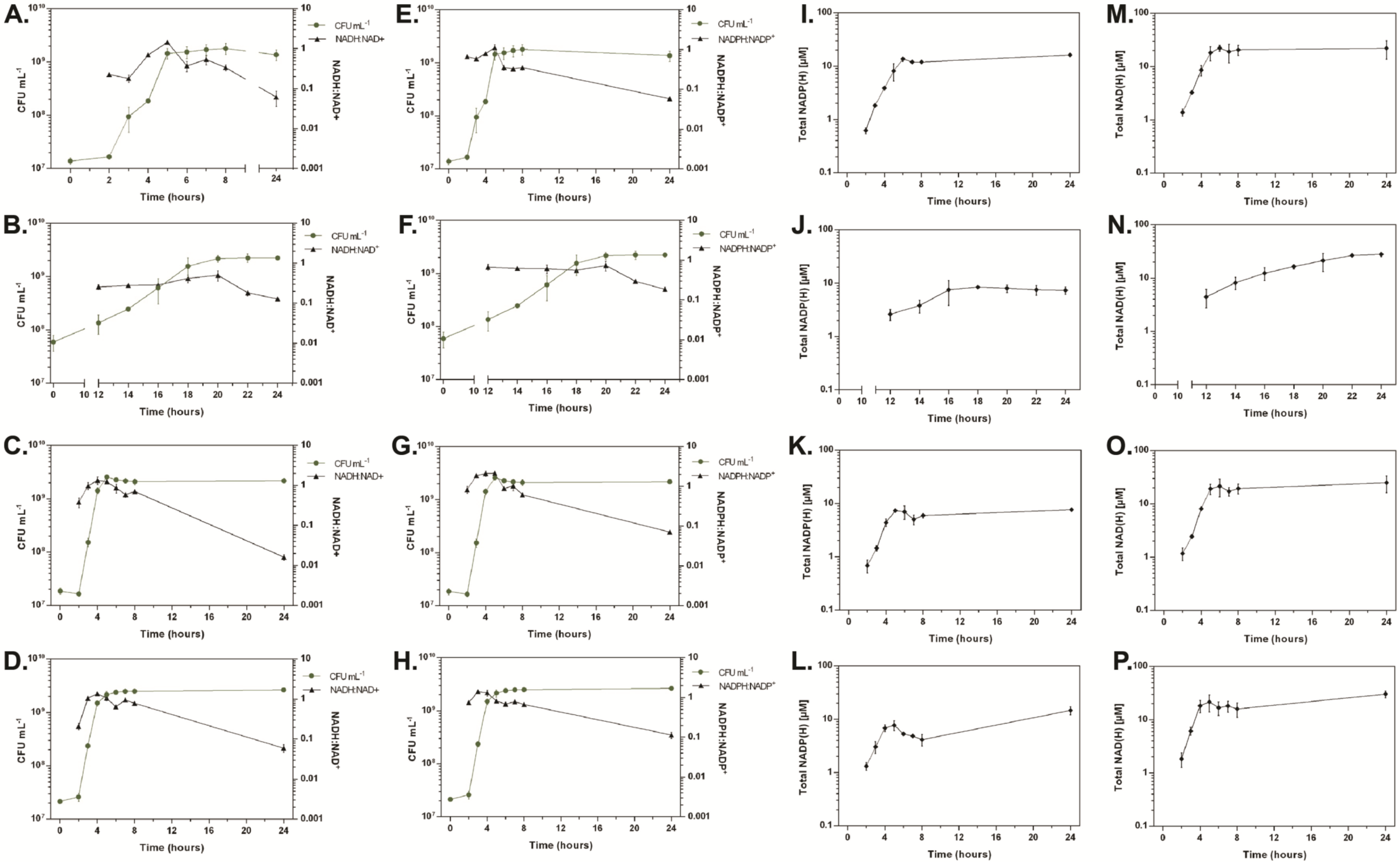

### Figure_S3

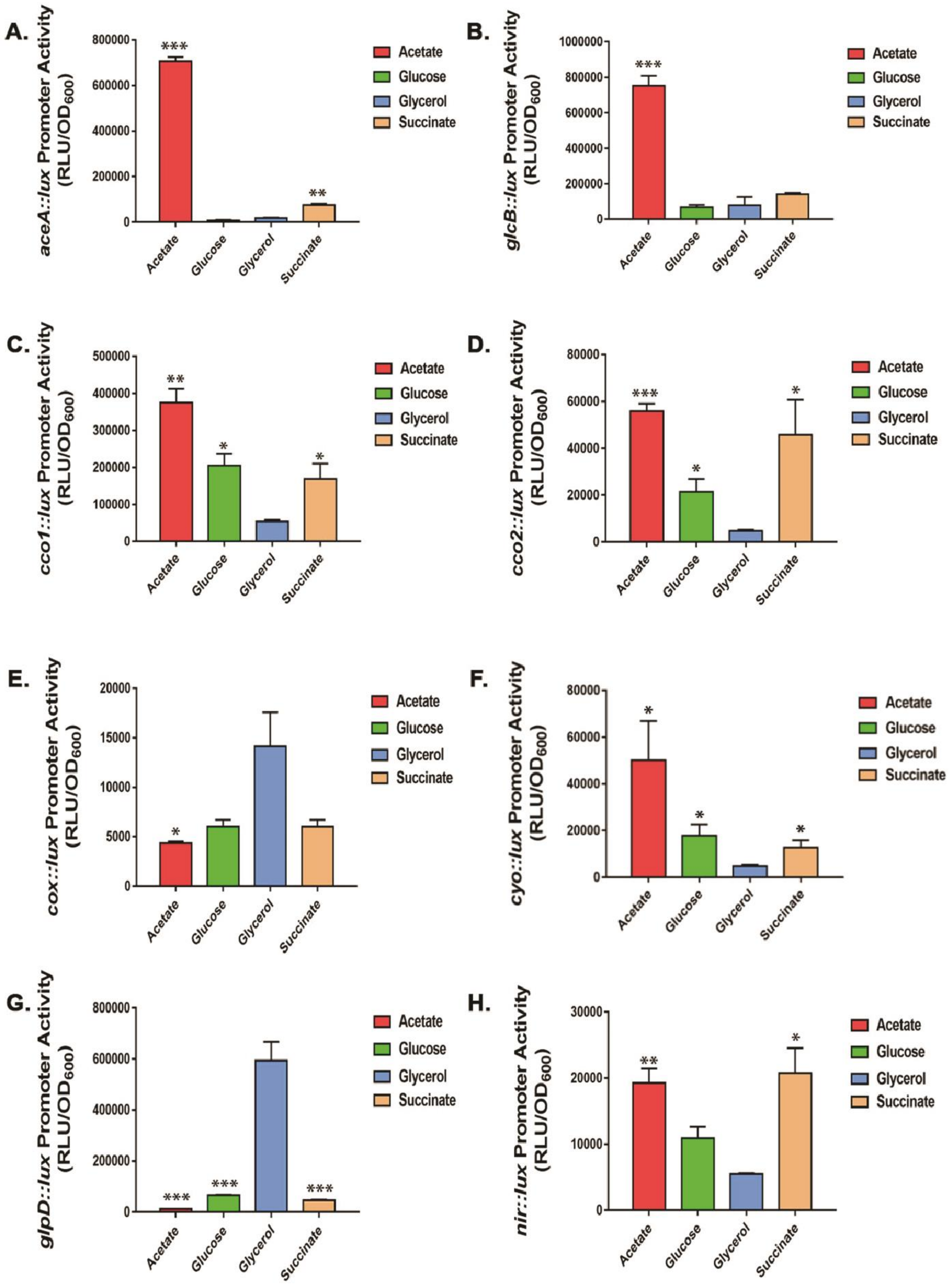

### Figure_S4

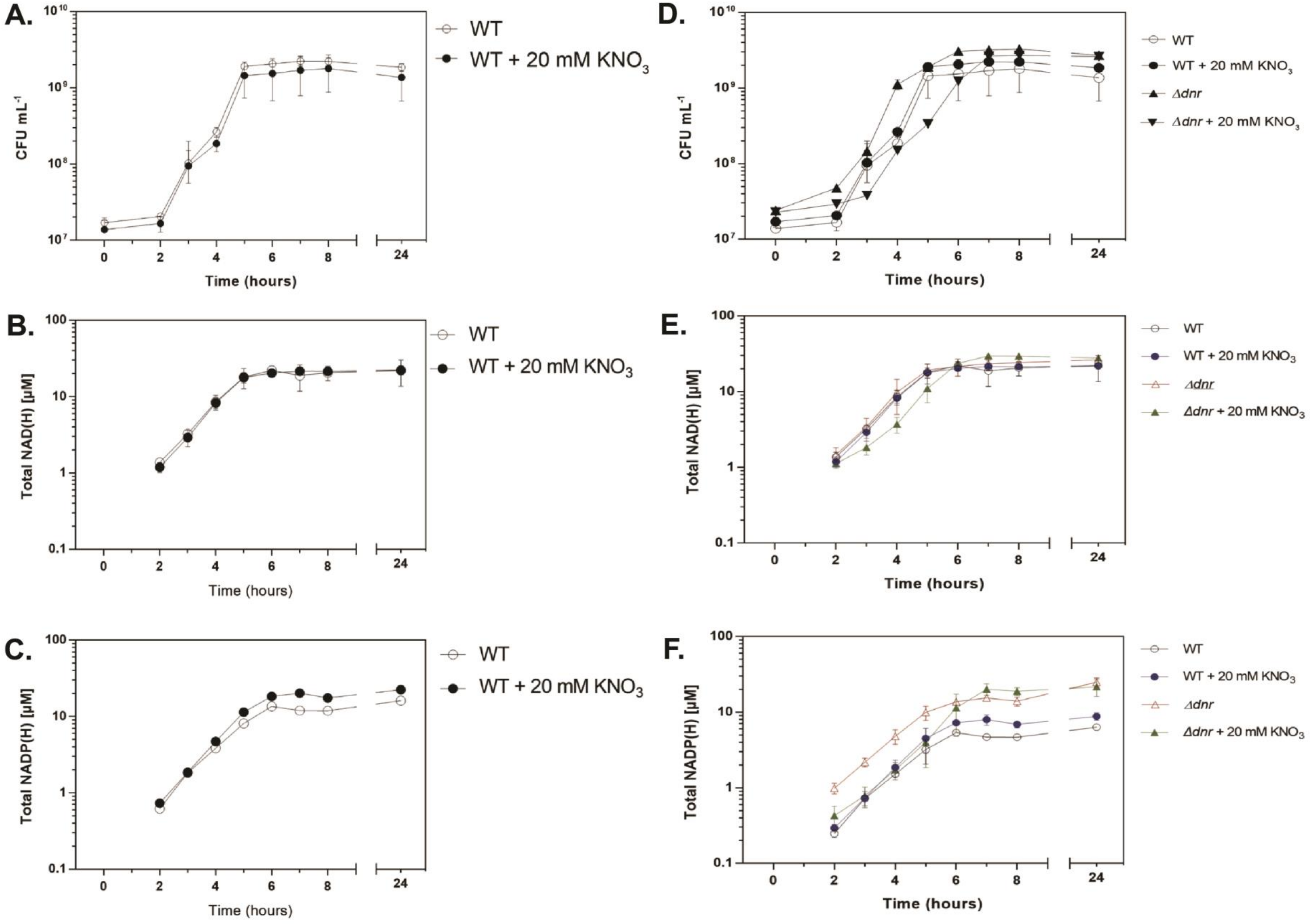

### Figure_S5

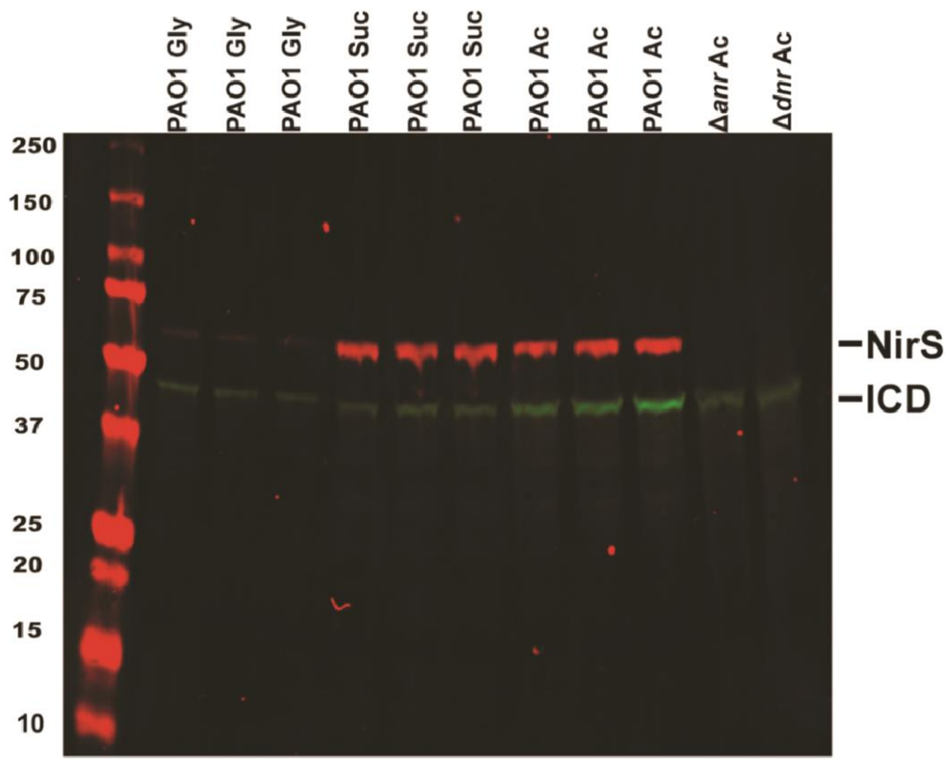

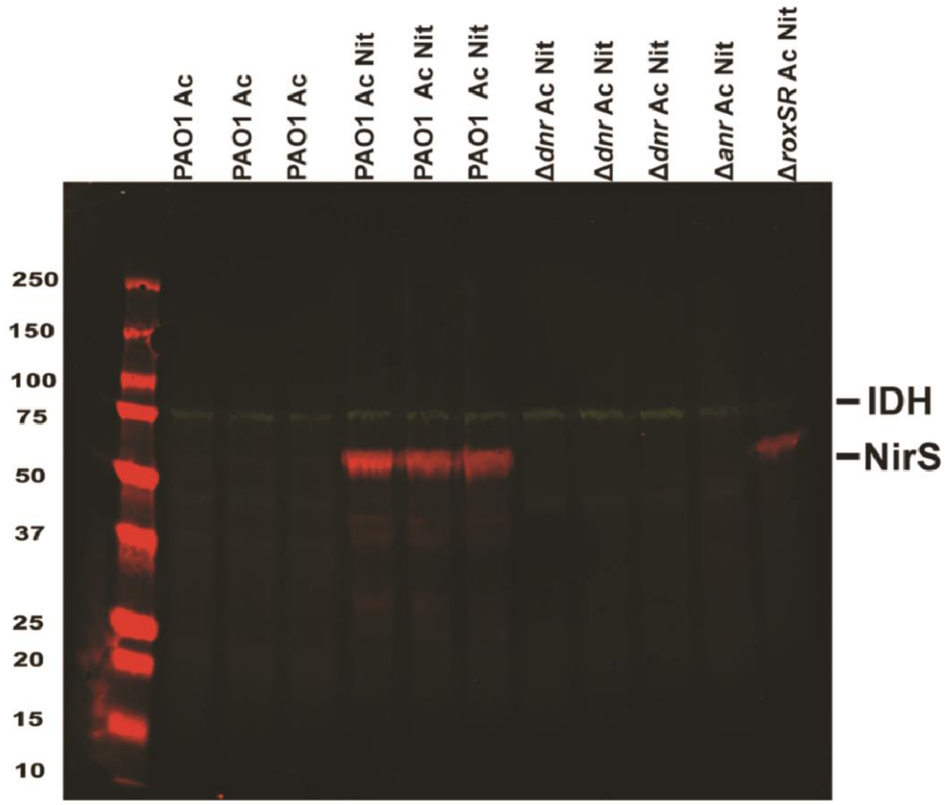

### Figure_S6

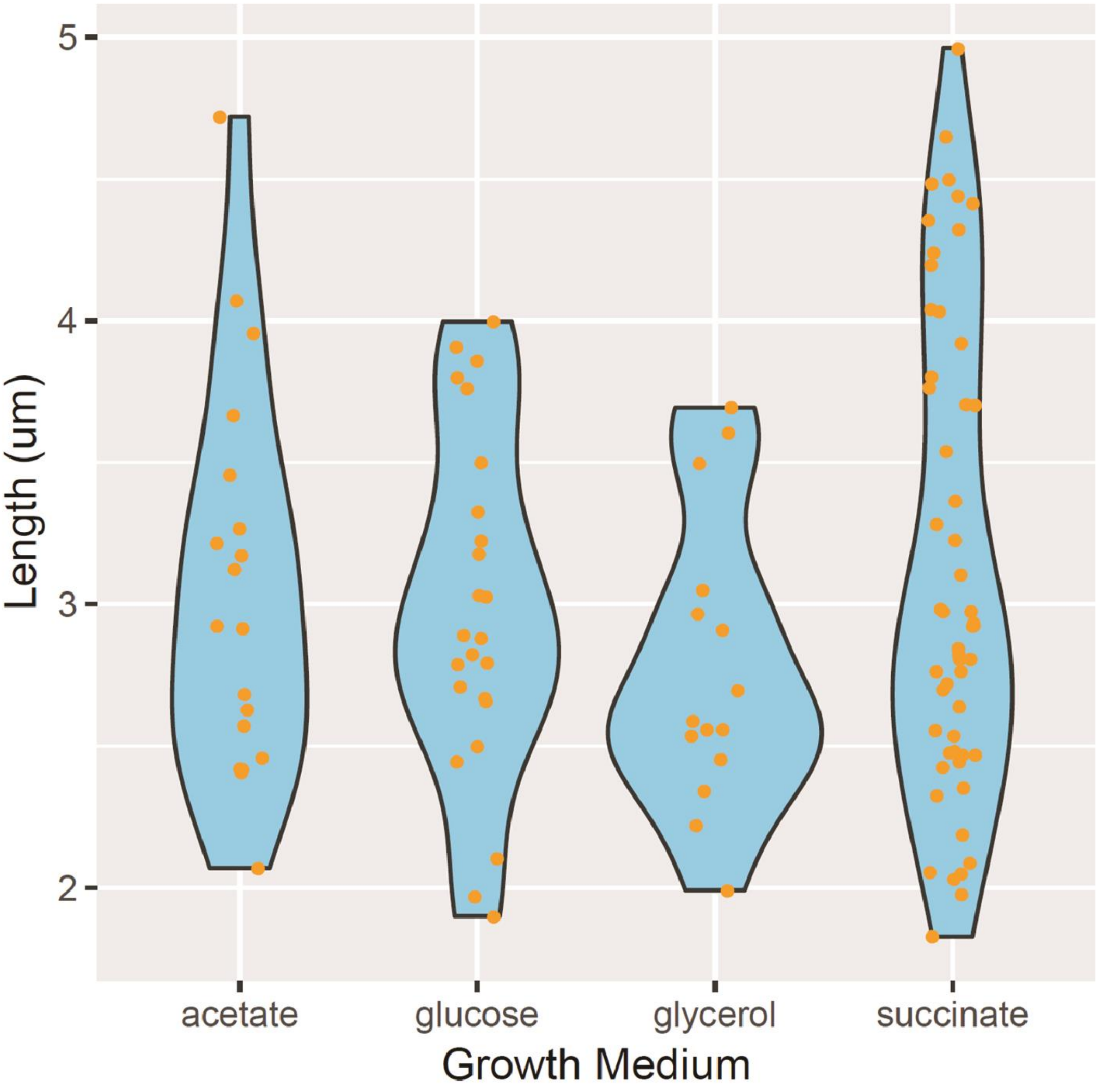
