## Supplementary material for "Contextual flexibility in *Pseudomonas aeruginosa* central carbon metabolism during growth in single carbon sources": Table_S1AB

**Table S1A.** Oligonucleotide primers used in this study.

| **Primer Name** | **Sequence (5′ to 3′)** |
|---|---|
| *ANR* KO UP F | AcccggggatcctctAGTTACTTCGCCGAGCTGTG |
| *ANR* KO UP R | TgcacttcGGCCAGACTGCAATCCTTG |
| *ANR* KO DW F | GtctggccGAAGTGCACATCCTCGAC |
| *ANR* KO DW R | CtgcaggtcgactctCTGAAGCTGAAATCCATC |
| *DNR* KO UP F | AcccggggatcctctGTACATGGTCGAACGCCTG |
| *DNR* KO UP R | aggcgttcCAGGTGGTGGCTTTGCAG |
| *DNR* KO DW F | AccacctgGAACGCCTGGAGTGCTTC |
| *DNR* KO DW R | CtgcaggtcgactctATGCCTGGCTCGACTTCC |
| *roxSR* KO UP F | acccggggatcctctATCTGCACCTCGATCACAC |
| *roxSR* KO UP R | cgtcgatgGGAGAGCAGGAAAACCGG |
| *roxSR* KO DW F | tgctctccCATCGACGGACCTTGCAG |
| *roxSR* KO DW R | ctgcaggtcgactctGTACGAGGGGATGCTTCAG |
| *pEX19Gm* F | AGAGTCGACCTGCAGGCATG |
| *pEX19Gm* R | AGAGGATCCCCGGGTACC |
| *aceA Tn7T lux* F *BamHI* | AAACGCGGATCCCAGCGAACAGAACCAGGC |
| *aceA Tn7T lux* R *XhoI* | AAAACTCGAGGCTGCCGTTCTTCTCTTTCA |
| *glcB Tn7T lux* F *BamHI* | AAACGCGGATCCGTAGAAGTCGAGGTAGGCGG |
| *glcB* Tn7T lux R *XhoI* | AAAACTCGAGTCCAGAACGTGTCGGCAGs |

| | |
|---|---|
| *cco1 Tn7T lux* F *BamHI* | AAACGCGGATCCCCCAGCTCCAACAAACCATC |
| *cco1 Tn7T lux* R *XhoI* | AAAACTCGAGGACACCGAGACCCATTCCAA |
| *cox Tn7T lux* F *BamHI* | AAACGCGGATCCTGAGTTCACGGAGGGCAG |
| *cox Tn7T lux* R *XhoI* | AAAACTCGAGCGAGAGCAAAAGGAAGCCC |
| *cco2 Tn7T lux* F *BamHI* | AAACGCGGATCCCCATGTAGGGAAACTCGAAGC |
| *cco2 Tn7T lux* R *XhoI* | AAAACTCGAGACGGTCATGATGGCGAATTG |
| *glpD Tn7T lux* F *BamHI* | AAACGCGGATCCTCAACCGGGTCATCAGCG |
| *glpD Tn7T lux* R *XhoI* | AAAACTCGAGCAAAGGAACACGGACAGGC |
| *dnr Tn7T lux* F *BamHI* | AAACGCGGATCCTCTATCCTGACATCCGTGCT |
| *dnr Tn7T lux* R *XhoI* | AAAACTCGAGCGAACAGGTGGTGGCTTTG |
| *cyo Tn7T lux* F *BamHI* | AAACGCGGATCCCGGATACAGTTGGCGCATCT |
| *cyo Tn7T lux* R *XhoI* | AAAACTCGAGTCGGGTTGAACAGGGTCATG |
| *nir Tn7T lux* F *BamHI* | AAACGCGGATCCCATGTACTGGACGAAGCGG |
| *nir Tn7T lux* R *XhoI* | AAAACTCGAGCGGCTTTCATGTCGTCCTTG |

**Table S1B:** Bacterial strains and plasmids used in this study

| Strain or plasmid | Description | Source or references |
| --- | --- | --- |
| **Strains** | | |
| *E. coli* JM109 | *E. coli* strain for cloning and expression | New England Biolabs |
| *P. aeruginosa* PAO1 | *P. aeruginosa* reference isolate | (1) |
| PAO1 *Δanr* | Anr is a transcriptional activator of anaerobic gene expression. | This study |
| PAO1 *Δdnr* | Dnr is a transcriptional activator of denitrification gene expression. | This study |
| PAO1 *ΔroxSR* | RoxSR is a redox-responsive two-component transcriptional regulator | This study |
| **Plasmids** | | |
| pEX19Gm | *P. aeruginosa* suicide vector, Gm | (2) |
| pUC18T-mini-Tn7T-lux-Gm | mini-Tn7 *luxCDABE* transcriptional fusion vector | (3) |
| pMF230 | Broad host-range plasmid for constitutive expression of the eGFP. Developed for imaging *P. aeruginosa*. | (4) |
